# nPhase: An accurate and contiguous phasing method for polyploids

**DOI:** 10.1101/2020.07.24.219105

**Authors:** Omar Abou Saada, Andreas Tsouris, Anne Friedrich, Joseph Schacherer

## Abstract

While genome sequencing and assembly are now routine, we still do not have a full and precise picture of polyploid genomes. Phasing these genomes, *i.e.* deducing haplotypes from genomic data, remains a challenge. Despite numerous attempts, no existing polyploid phasing method provides accurate and contiguous haplotype predictions. To address this need, we developed nPhase, a ploidy agnostic pipeline and algorithm that leverage the accuracy of short reads and the length of long reads to solve reference alignment-based phasing for samples of unspecified ploidy (https://github.com/nPhasePipeline/nPhase). nPhase was validated on virtually constructed polyploid genomes of the model species *Saccharomyces cerevisiae*, generated by combining sequencing data of homozygous isolates. nPhase obtained on average >95% accuracy and a contiguous 1.25 haplotigs per haplotype to cover >90% of each chromosome (heterozygosity rate ≥0.5%). This new phasing method opens the door to explore polyploid genomes through applications such as population genomics and hybrid studies.

## Introduction

Studying genotype-phenotype relations is contingent on having an accurate view of the genetic variants. To that end, various sequencing strategies and ways to analyze them have been developed. The ultimate goal is to faithfully determine the precise sequence of the DNA molecules contained within the cell. In practice this level of precision is rarely necessary and approximations are routinely used when they can be afforded. Aligning the sequencing data to a reference genome is a good approximation to identify genetic variants such as Single Nucleotide Polymorphisms (SNPs) but a poor one to identify Structural Variants (SVs)^1^. By contrast, the generation of *de novo* assemblies using the sequencing data is a good approximation to identify SVs^1^ but, without significant polishing work^2^, usually leads to a lower quality sequence. One enduring approximation is the reduction of the genome to a single sequence, even if the organism does not have a haploid or rigorously homozygous genome. A diploid or higher ploidy genome can be heterozygous. Identifying the heterozygous positions, or variants, is known as genotyping. Linking these variants together to establish which variants co-occur on the same strand of DNA is known as haplotyping or phasing. There is increasing interest in phasing genomes for diverse reasons, such as to obtain more accurate reference genomes^3^, better study population genomics^4^, improve the accuracy of GWAS studies^5^, study the effects of compound heterozygosity^6^, investigate Allele-Specific Expression patterns^7^, gain insight into polyploid evolution^8,9^, better understand the mechanisms of heterosis^10^ and dissect the origins of hybrid species^11^.

Phased genomes can be obtained either by physically separating entire chromosomes^12^ (or significantly large portions of chromosomes) prior to sequencing^13^ or by separating them bioinformatically after sequencing the whole genome^14^. The length of reads is a significant limiting factor in the ability to bioinformatically separate reads into their corresponding haplotypes. One very successful method that overcame that limitation was trio binning^15^, which circumvented the importance of long reads by leveraging information from parental whole genome sequencing. Other methods have been explored but cannot overcome the short read length limitation particularly well^16^. One solution has been to resort to imputing haplotypes through reference panels^17^. Despite a higher error rate, diploid phasing of long reads has been solved by existing methods such as WhatsHap^18^, an alignment-based phasing tool and Falcon-Unzip^19^, an assembly-based phasing tool. Assembly-based phasing attempts to generate a *de novo* assembly for each haplotype directly, without relying on a reference sequence. Alignment-based phasing uses a reference genome as support to identify heterozygous positions and then attempts to link positions together based on the co-occurrence of heterozygous SNPs on overlapping reads. For diploids each variable position can only be one of two possible bases. Knowing one haplotype allows to deduce the other. This allows diploid phasing methods to be relatively simple and straight-forward. For polyploids, however, a variable position can be one of two or up to six possible states (all four bases, a deletion or an insertion) and this deduction is no longer possible, rendering the task of phasing significantly more complex. Some methods currently exist to phase polyploids but mainly using short read sequencing and leading to a low accuracy and contiguity phasing^20,21,22,23^.

Here, we developed nPhase to address the lack of a polyploid phasing method that outputs accurate, contiguous results and does not require prior knowledge of the ploidy of the sequenced genome. The required inputs are a reference sequence as well as long and short read sequencing data. The pipeline performs the mapping, variant calling, phasing and outputs the phased variants and a fastQ file for each predicted phased haplotype, or haplotig. The nPhase algorithm is ploidy agnostic, meaning it does not require any prior knowledge of ploidy and will not attempt to guess the ploidy of the sample. Instead it will separate the reads into as few distinct haplotigs as possible. The nPhase algorithm has three modifiable parameters, we have evaluated the effects of these parameters on the results and provide a default set of parameters, which we predict to be appropriate for all cases, along with some recommendations on how to modify these parameters for genomes that are more difficult to phase, *i.e.* low heterozygosity and high ploidy genomes.

Using the yeast species *Saccharomyces cerevisiae* as a model, we validated the performance of nPhase on simulated genomes (2n, 3n and 4n) of varying heterozygosity levels (0.01%, 0.05%, 0.1% and 0.5% of the genome). We found that nPhase performs very well in terms of accuracy and contiguity. We obtained an average of 93.9% accuracy for all diploids, 92.3% for all triploids, and 94.5% for tetraploids with a heterozygosity level of at least 0.5%, or 87.3% accuracy when we include the lowest heterozygosity level tetraploids. All results are very contiguous, with an average of between 2.4 and 4.1 haplotigs per haplotype, bringing us very close to the ideal result of one haplotig per haplotype.

## Results

### Phasing pipeline and strategy

We developed the nPhase pipeline, an alignment-based phasing method and associated algorithm that run using three inputs: highly accurate short reads, informative long reads and a reference sequence. The pipeline takes the raw inputs and processes them into data usable by the nPhase algorithm. Unlike other existing methods, our algorithm is designed for ploidy agnostic phasing. It does not require the user to input a ploidy level and it does not contain any logic that attempts to estimate the ploidy of the input data. The idea at the core of the algorithm is that if you iteratively cluster the most similar long reads and groups of long reads together you will naturally recreate the original haplotypes.

For the first step of the pipeline, the long and short reads are aligned to the reference, then the aligned short reads are variant called to identify heterozygous positions and generate a high quality dataset of variable positions (Figure 1). Each long read is then reduced to its set of heterozygous SNPs according to the previously identified variable positions. We also collect long read coverage information to allow the level of representation of a haplotype in the data to influence its likelihood of being properly phased (see Methods).

**Figure 1.**
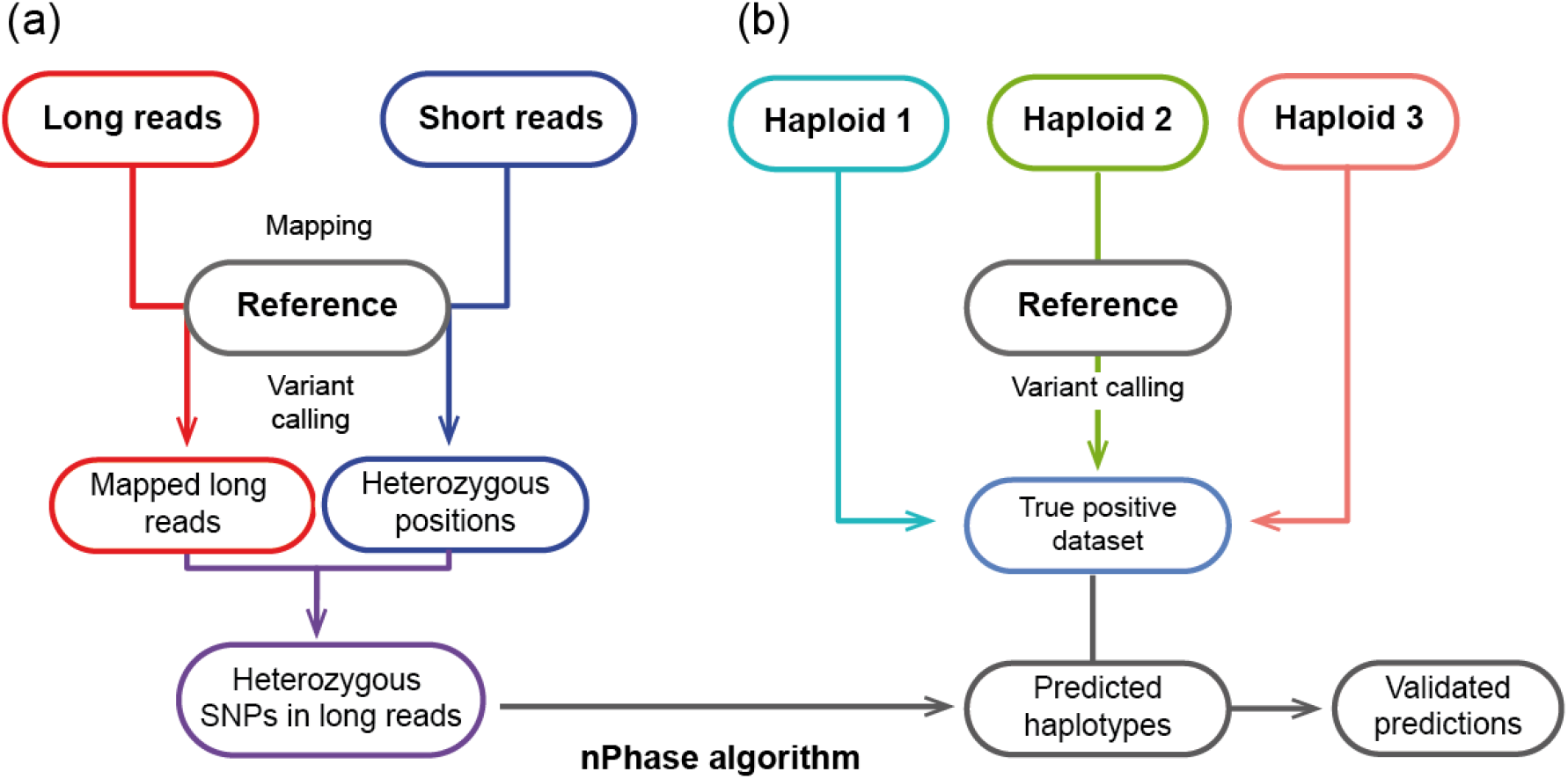
nPhase pipeline and verification process. (**a**) The nPhase pipeline requires three inputs: a long read dataset, a short read dataset and a reference genome sequence. Both sequencing datasets are mapped to this reference genome, then the short reads are variant called in order to identify heterozygous positions. The long reads are reduced to only their heterozygous positions, and this set of linked heterozygous positions is phased by the nPhase algorithm and outputs phased haplotypes. (**b**) In parallel with running the virtual polyploids through the nPhase pipeline, we map the original strains to the same reference and variant call them to identify their haplotypes. This generates the true positive dataset against which we will compare the haplotypes predicted by nPhase in order to assess the accuracy of our algorithm.

The reduced long reads and the coverage information are then passed onto the nPhase algorithm, an iterative clustering method. On the first iteration, nPhase clusters together the two most similar long reads, then it checks that the cluster identities are maintained, *i.e.* it checks that merging these two long reads together does not significantly change the information they each contain individually, and finally it generates a consensus sequence representative of the group of these two reads. The next iteration will be exactly the same with N-1 reads. nPhase will run until all remaining clusters are sufficiently different from each other to fail the cluster identity maintenance check. These remaining clusters represent the different haplotypes within the original dataset.

### nPhase, a ploidy agnostic phasing algorithm

As described earlier, nPhase is an iterative clustering algorithm. It is composed of three main ideas: (i) clustering, which ensures that similar reads are clustered together, (ii) cluster identity maintenance, which ensures that only similar clusters are merged into larger ones and finally (iii) consensus, a way to reduce a cluster to a consensus sequence in order to easily compare it to other clusters (Figure 2).

**Figure 2.**
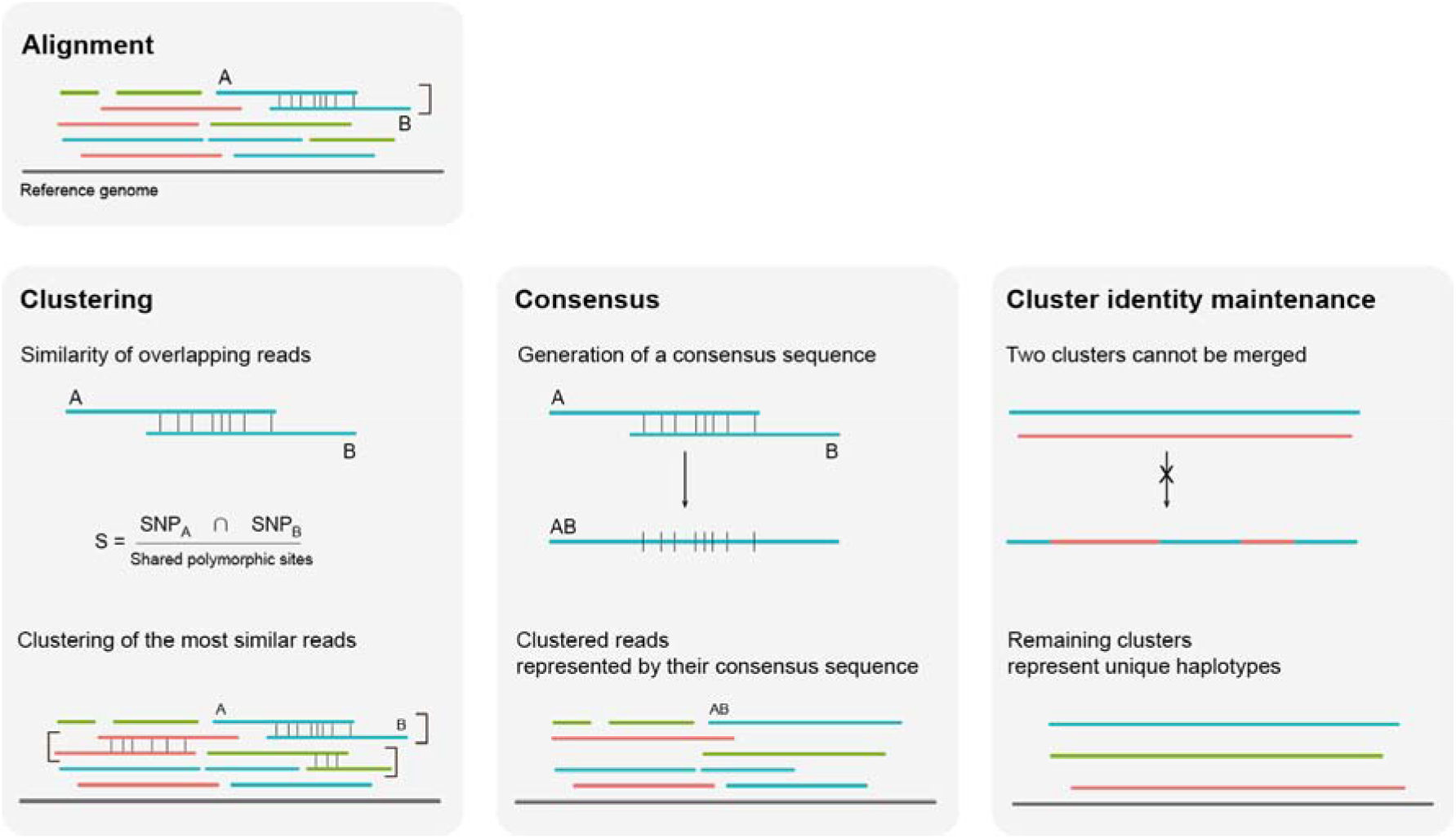
nPhase algorithm. Here we represent how a triploid’s reads could align to a reference sequence. Each read is one of three colors, one for each haplotype. The clustering, consensus and cluster identity maintenance steps are iteratively repeated until all remaining clusters are forbidden to merge. **Clustering**: Each vertical line represents a SNP; different colors signify different haplotypic origins. Only two reads are clustered at a time, here we show three clusters, so this is the result of the third step of nPhase’s iterative clustering. **Consensus**: A consensus sequence is generated by allowing every read in the cluster to vote for a specific base for a given position. Votes are weighted by the pre-calculated context coverage number to discourage sequencing errors. The consensus sequences that represent clusters are treated just like aligned long reads and continue to be clustered. **Cluster identity maintenance**: When all remaining clusters are very different from each other they are not allowed to merge, this is to prevent the algorithm from always outputting only one cluster per region. The remaining clusters and their consensus sequences should correspond to the haplotypes present in the original dataset.

Each step of the clustering algorithm starts by calculating the similarity between every overlapping pair of reads (Figure 2a). By default, the minimal overlap is 10 heterozygous positions. Similarity is defined as S = N_shared variants_/N_shared positions_. The pair of reads with the highest similarity is clustered together. If there is a tie, then we cluster together the pair of reads with the most variable positions in common. If there is again a tie, then we select a pair randomly. By default, the algorithm will not attempt to merge two sequences with less than 1% similarity.

The pair that was selected now forms a cluster of two reads (Figure 2b). In order to continue this iterative algorithm, we need to define a way to calculate the similarity between a read and a cluster of reads, and the similarity between two clusters of reads. We do so by computing a consensus sequence for each cluster of reads and we use the consensus sequence to calculate the similarity as defined above. For each position, the consensus is defined as the base which has the most support from the reads in the cluster. Each read gets a vote equal to the context coverage of the base it supports. If there is a tie then all tied bases are included in the consensus sequence.

As defined, the clustering algorithm will continue to iterate, merging clusters together until all available options are exhausted and output only one cluster per region (Figure 2c). The solution is to set restrictions on which clusters are allowed to be merged in the clustering step. We consider that each cluster has its own “identity” defined by the population of reads that comprise it. If merging two clusters has a significant effect on the identity of both clusters then the merge is not allowed. We calculate how much merging of two clusters would change them. The amount of change allowed is limited by the ID parameter. In order to quantify the amount of change to a cluster’s identity we keep track of the “demographics” of each position, *i.e.* how strongly represented each base is for that position in that cluster. We differentiate positive identity changes from negative identity changes: (i) if a merge of two clusters results in increased support for their consensus sequence bases then that change is considered positive, (ii) if the merge results in decreased support for a consensus sequence base then that change is considered negative and we calculate how many votes the base lost, even if it remains the consensus base after the merge. The number of votes lost is divided by the total number of votes in the region that both clusters have in common to obtain the cluster identity change percentage. By default, if it is higher than 5% we do not allow the two clusters to merge. Once all remaining clusters fail this test, the algorithm stops. The resulting clusters represent the different haplotypes that nPhase found and are output as different sets of reads, heterozygous SNPs, and consensus sequences.

### Validation of the nPhase algorithm by combining reads of non-heterozygous individuals

To test and validate the performance of nPhase, we decided to combine sequencing datasets of haploid and homozygous diploid organisms into virtual polyploid datasets. We selected four natural *S. cerevisiae* isolates as the basis for our virtual genomes: ACA, a haploid strain, and three homozygous diploid strains: CCN, BMB and CRL (Supplemental Table 1). These four strains have different ecological and geographical origins and are sufficiently distinct from each other to allow us to evaluate the performance of nPhase at heterozygosity levels of up to 1% of the genome^24^.

We sequenced these strains using an Oxford Nanopore long-read sequencing strategy and obtained Illumina short-read data from our 1,011 yeast genomes project^24^. Since these strains do not have any heterozygosity, we could map their short reads to the *Saccharomyces cerevisiae* reference genome and variant call them to obtain their haplotypes (Figure 1). We then used these haplotypes as a truth set to assess the performance of nPhase. With this truth set, we tested the influence of dataset characteristics: coverage, ploidy, heterozygosity level and the inclusion or exclusion of reads that map to distant regions of the genome, hereafter described as split reads. We also investigated the influence of parameters that modulate the behavior of the nPhase algorithm: minimum similarity, minimum overlap and maximum ID change (for a description of them see **Available Parameters** in Methods).

To assess the influence of ploidy, we used the three constructions of the different virtual genomes previously mentioned. We also randomly sampled 6250, 12500, 62500 and 125000 heterozygous SNPs from each virtual genome to simulate datasets where 0.05%, 0.1%, 0.5% and 1% of the positions in the genome are heterozygous. This equates to three different ploidies and four heterozygosity levels, or 12 polyploid genomes to test.

By running a total of 6000 validation tests on varying ploidy, heterozygosity, and coverage levels exploring the parameter space, we determined default parameters of nPhase (see Methods). According to these tests, the parameters that result in optimal results in terms of accuracy and contiguity are the following: 1% minimum similarity, 10% minimum overlap and 5% maximum ID (see **Identifying optimal parameters** in Methods). We then ran nPhase with these default parameters on our previously described optimal datasets of varying ploidy (2n, 3n and 4n) and heterozygosity levels (0.05%, 0.1%, 0.5% and 1%) of 20X long reads per haplotype with split read information (Supplemental Table 2).

As an example, we phased the tetraploid genome showing a heterozygosity level of 0.5% using nPhase (Figure 3). Since we know the ground truth, we can assign each haplotig to the strain whose haplotype it most closely represents and we can calculate our accuracy metrics.

**Figure 3.**
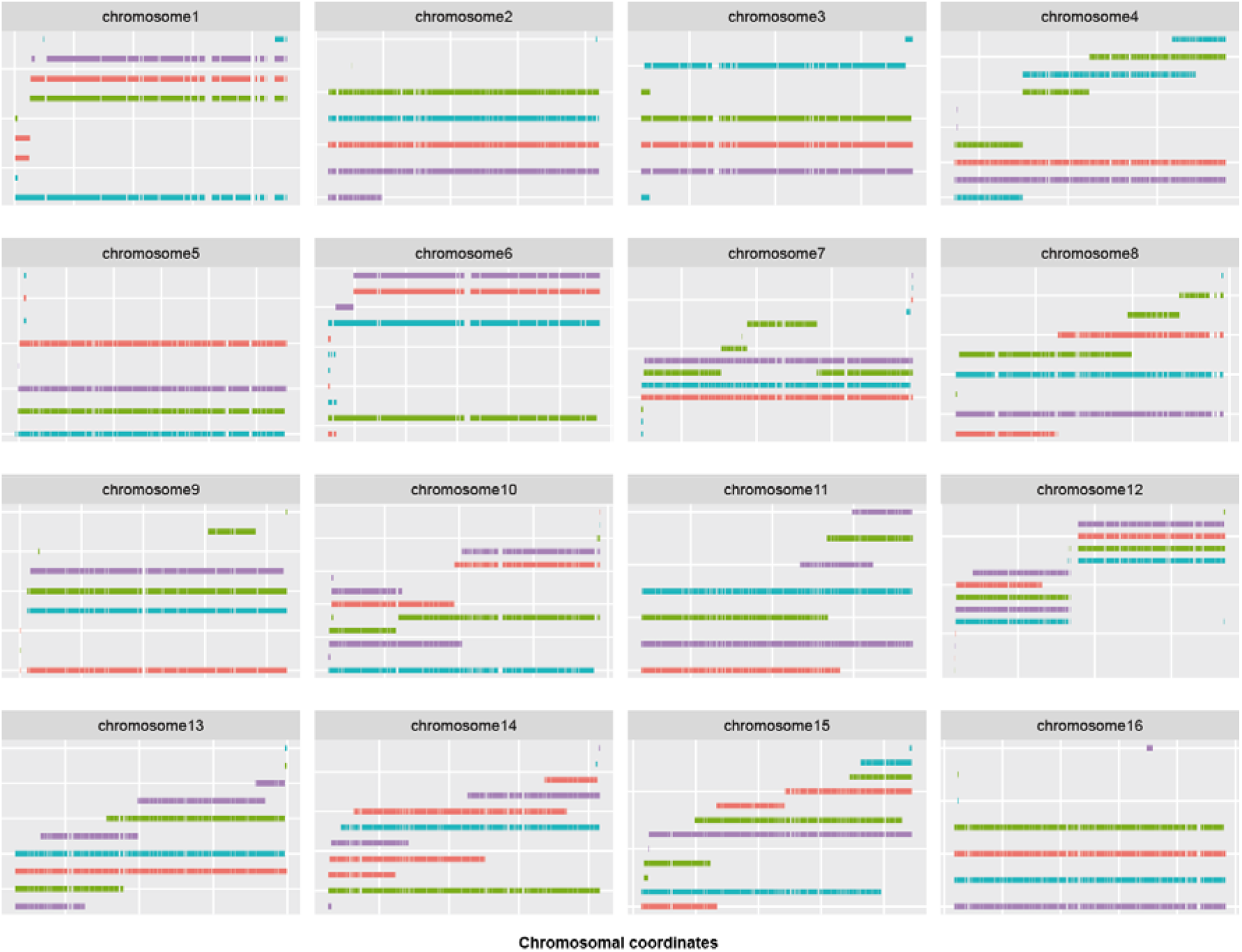
Predicted haplotypes for the tetraploid genome with a 0.5% heterozygosity level. The result of this test was an accuracy of 93.7%, an error rate of 4.0%, and a missing rate of 2.2% with an average of 2.4 haplotigs per haplotype. Each subgraph displays the predicted haplotigs for a different chromosome, each predicted haplotig is on a different row on the Y axis, and the X axis displays the position along the chromosome. All predicted haplotigs are color coded according to the haplotype they are the closest to.

In order to measure accuracy we distinguish between two forms of errors: standard errors, *i.e.* heterozygous SNPs erroneously attributed to the wrong haplotype, and missing errors, *i.e.* heterozygous SNPs which we know are present but which were erroneously not represented in the predictions. The accuracy is the percentage of all SNPs which were correctly attributed to their haplotype. The error rate is the percentage of all predictions which were incorrect. The missing rate is the percentage of all heterozygous SNPs which were never attributed to their haplotype. We use the following formula: accuracy=TP/(TP+FP+FN) with

TP=True Positive; the SNP was attributed to the correct haplotype

FP=False Positive; the SNP does not belong in this haplotype

FN=False Negative; the SNP is not represented in the results.

The result of this test was an accuracy of 93.7%, an error rate of 4.0%, and a missing rate of 2.2% with an average of 2.4 haplotigs per haplotype. Seven of the sixteen chromosomes have an L90 of 1, meaning that for all four haplotypes, more than 90% of the heterozygous SNPs were assigned to one haplotig. For the nine remaining chromosomes, seven have at least two chromosome-length haplotigs. In all cases, the chromosomes are nearly fully covered by haplotigs that represent the four different haplotypes, as confirmed by the low missing haplotype prediction rate (2.2%). As a ploidy agnostic tool, nPhase was not given any information about the ploidy of this sample and does not attempt to estimate its ploidy. Despite that, nPhase reached a high accuracy (93.7%) and contiguity (2.4 haplotigs per haplotype), demonstrating its ability to reliably phase a tetraploid of that heterozygosity level. The same representation is available for the other datasets of different ploidy and heterozygosity levels (Supplemental figure 1).

Across the 12 phased genomes with variable ploidy and heterozygosity levels, we noted little variation in terms of contiguity as we obtained between 2.4 and 4.3 haplotigs per haplotype (Figure 4a). At a heterozygosity level of 0.05%, the least contiguous genomes are observed with around 4 haplotigs per haplotype (Figure 4a). The triploid genomes decrease to around 3 haplotigs per haplotype for heterozygosity levels greater than 0.1% (Figure 4a). The tetraploid tests continue the trend of higher ploidies becoming more stable and contiguous as the heterozygosity level increases, dropping to 3.1 haplotigs per haplotype at the 0.1% heterozygosity level and then stabilizing at 2.4 haplotigs per haplotype at the 0.5% and 1% heterozygosity levels (Figure 4a). This could be explained by the availability of more haplotigs to potentially merge with each other as ploidy increases.

**Figure 4.**
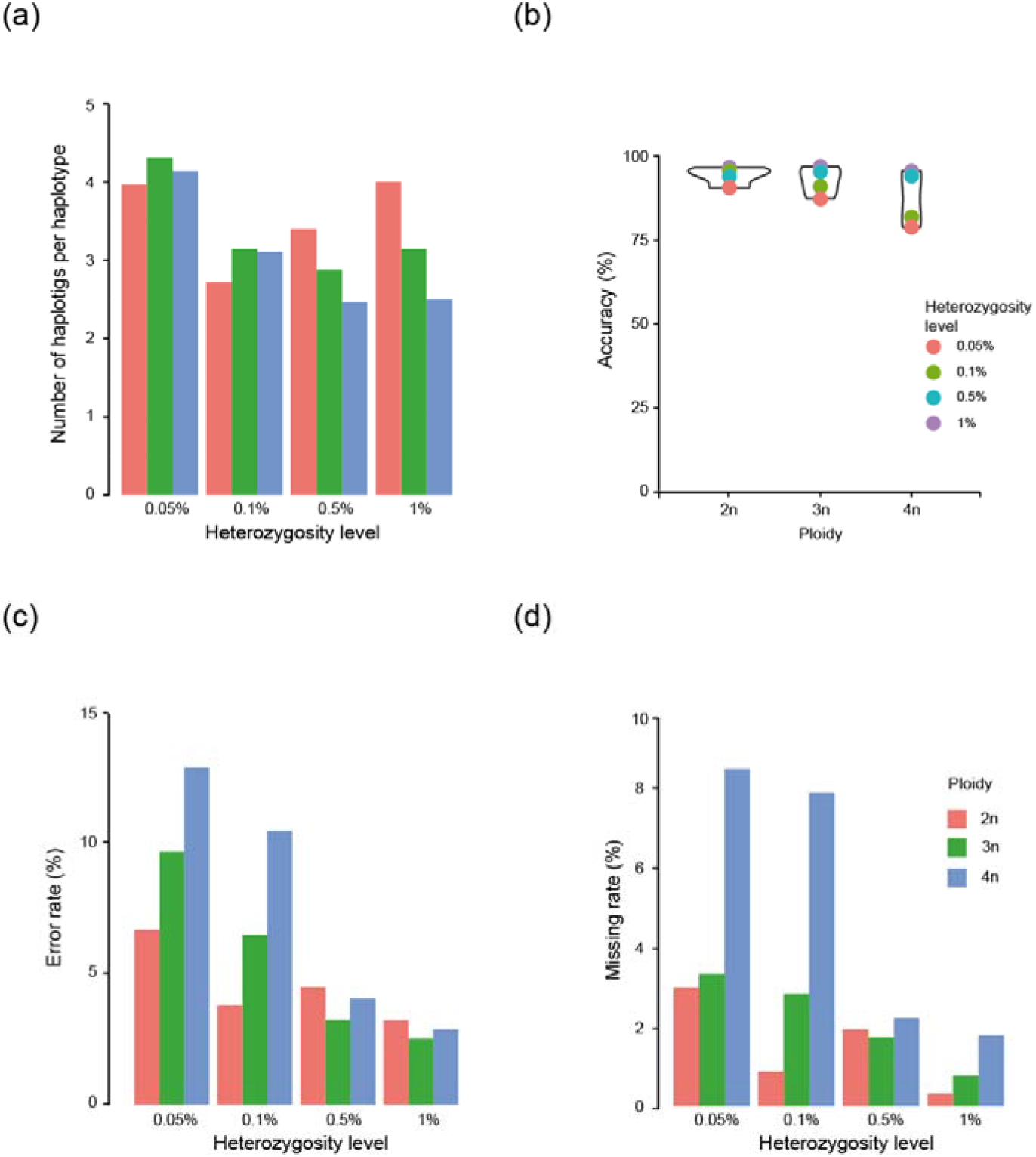
Effects of ploidy and heterozygosity levels on accuracy and contiguity. Through these graphs we show the effects of sample properties (heterozygosity level and ploidy) on nPhase’s accuracy metrics when run with default parameters. (**a**) Each bar displays the contiguity of a different test result. The least contiguous heterozygosity level is 0.05%, likely related to its also yielding the least accurate results. Overall, we note little absolute variation in the contiguity. Interestingly, contiguity at higher heterozygosity levels appears to be a function of ploidy. Higher ploidies seem less likely to become less contiguous as a result of increasing the heterozygosity level, while the diploid tests are more affected. We also note that tetraploids of high heterozygosity level are the most contiguous. (**b**) Each bar displays the accuracy of a different test result. As ploidy increases, the accuracy tends to decrease. It also appears to decrease faster for tests on low heterozygosity level constructions. (**c** and **d**) Each bar displays our evaluation of the effects of ploidy and heterozygosity level on the error and missing rates, respectively, for our 12 tests using optimal parameters. Overall, we see that the error rate is always higher than the missing rate across these conditions. As the heterozygosity level increases, the error and missing rates decrease along with the gap between ploidies. We also find that more difficult phasing problems (high ploidy and low heterozygosity level) yield much higher error and missing rates, and that the low heterozygosity tetraploids seem to be particularly sensitive to missing calls.

Regarding the accuracy, we observed that for heterozygosity levels greater than 0.5%, the accuracy appears stable and high across ploidies with a minimum of 93.56% for the diploid (2n) at a 0.5% heterozygosity level, and a maximum of 96.70% accuracy for the triploid (3n) at a 1% heterozygosity level (Figure 4b). For lower heterozygosity levels (≤ 0.1%), we have results that are more variable between ploidies (Figure 4b). Diploid tests retain a high 95.32% accuracy for the 0.1% heterozygosity level but drop to 90.34% accuracy for the 0.05% heterozygosity level. For triploid genomes, the results drop to 90.70% accuracy for the 0.1% heterozygosity level, then down to 87.00% at 0.05% heterozygosity level. Continuing the trend of higher ploidies performing worse with lower heterozygosity levels, the accuracies for the 0.1% and 0.05% heterozygosity levels for the tetraploid tests output 81.65% and 78.62% accuracy, respectively.

In addition, we observed that errors are more frequent in all tests than missing calls (Figures 4c and 4d). For higher heterozygosity levels (≥ 0.5%), these two forms of error are stable and very low. The error rate is set between a minimum of 2.53% for the 1% heterozygosity level triploid and a maximum of 4.51% for the 0.5% heterozygosity level diploid. And the missing rate is set between a minimum of 0.31% for the 1% heterozygosity level diploid and a maximum of 2.21% for the 0.5% heterozygosity level tetraploid. For lower heterozygosity levels (≤ 0.1%), both the error and missing rates increase with ploidy, suggesting both types of errors may be linked. If we set aside the 0.1% heterozygosity level diploid which has an error and missing rates of 3.82% and 0.86%, respectively, the error rates have a wide range with a minimum error of 6.49% for the 0.1% heterozygosity level triploid and a maximum error of 12.91% for the 0.05% heterozygosity level tetraploid. Similarly, the missing rates range from a minimum of 2.97% for the 0.05% heterozygosity level diploid to a maximum of 8.46% for the 0.05% heterozygosity level tetraploid, again adding to the trend of lower heterozygosity levels coupled with higher ploidies yielding worse results.

### Benchmarking nPhase against other polyploid phasing tools

Some methods currently exist to phase polyploids using long read data such as WhatsHap polyphase^20^, as well as other methods which were mostly designed to work with short read sequencing data but can sometimes use long reads as input^21,22,23^. Because nPhase is a phasing tool that leverages the linking power of long reads to achieve its high accuracy and contiguity metrics, we did not benchmark it against tools that rely exclusively on short reads for phasing, since these are inherently limited by the size of their reads. We also did not benchmark nPhase against tools that can only phase diploid genomes as this is not the intended use case for our algorithm. We therefore compare nPhase to the recently released WhatsHap polyphase, to our knowledge the only other polyploid phasing algorithm that handles long reads.

We compared the results nPhase (default parameters) with WhatsHap polyphase on the same samples (Figure 5). Since WhatsHap polyphase has a parameter named “--block-cut-sensitivity” that can be set to determine the tradeoff between accuracy and contiguity, we tested WhatsHap polyphase using all possible values for this parameter (integers from 0 to 5) to compare all possible results to nPhase’s default results. A value of 0 for this parameter means that WhatsHap polyphase will generate the most contiguous results possible, and 5 means that it will generate the most accurate results possible.

**Figure 5.**
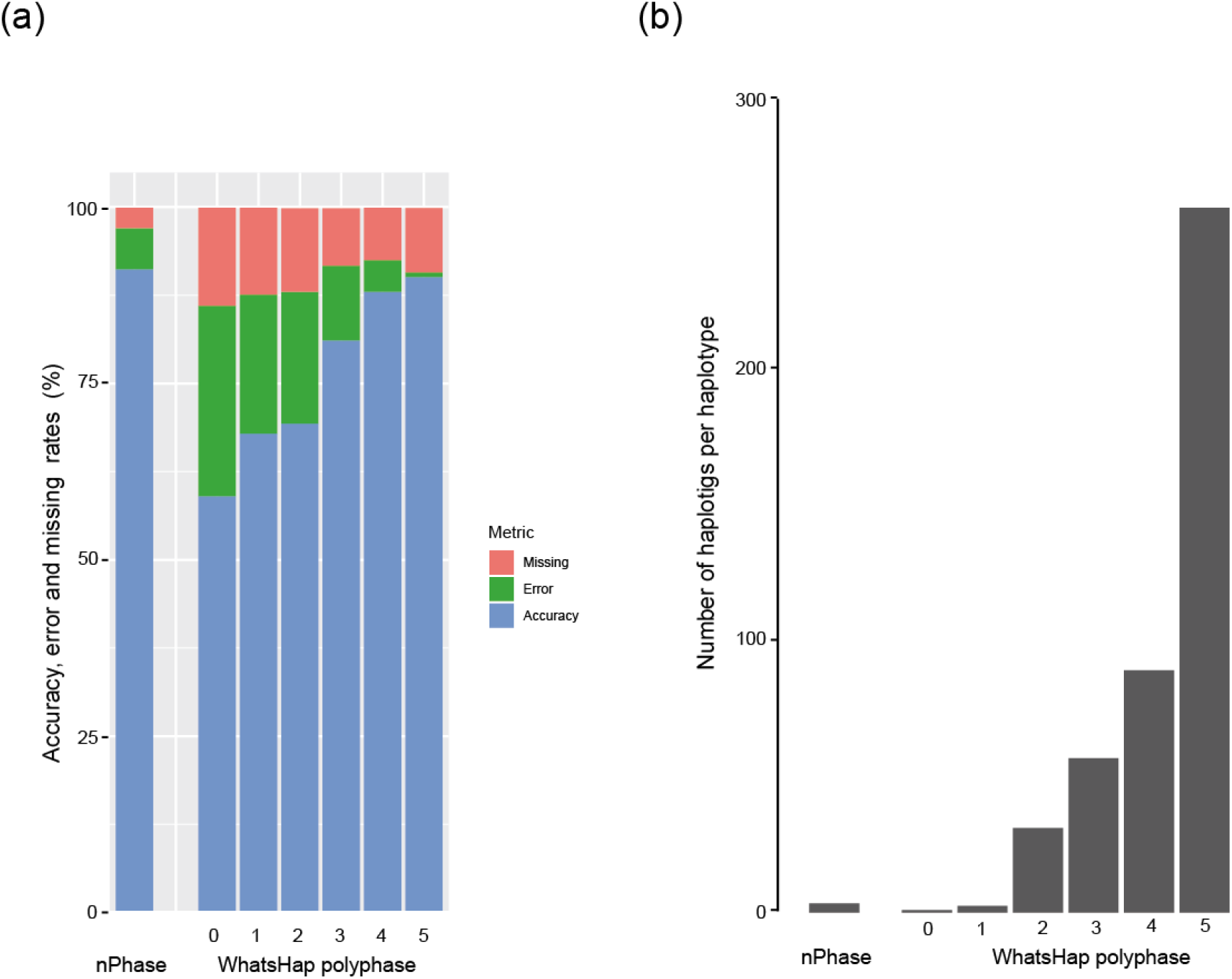
Error types and number of haplotigs for nPhase and WhatsHap polyphase. nPhase and WhatsHap polyphase were both applied to our 20X test datasets of different ploidy and heterozygosity levels. nPhase was tested using its default parameters and WhatsHap polyphase was tested with all six possible values of its adjustable sensitivity parameter. This graph compares both tools using the following metrics: average number of haplotigs obtained for the genome, normalized by the ploidy, average accuracy, average error rate and average missing rate. (**a**) Average accuracy, error and missing rates for all tests using nPhase and WhatsHap polyphase on different sensitivity levels. The error rate for WhatsHap polyphase increases dramatically as the sensitivity level decreases, illustrating the tool’s tradeoff between accuracy and contiguity. (**b**) Average number of haplotigs per chromosome per haplotype for all tests using nPhase and WhatsHap polyphase on different sensitivity levels. The very high number of haplotigs per chromosome per haplotype for the highest sensitivity levels (5 and 4) shows that despite being highly accurate, they are not contiguous enough to be informative. Based on our results, nPhase outperforms WhatsHap polyphase in all of our tests. The tradeoff between accuracy and contiguity is extreme in WhatsHap polyphase, either the results are very accurate but so fragmented as to be uninformative, or they are about as contiguous as nPhase but less than 60% accurate.

The performance of WhatsHap polyphase was measured in terms of switch error rate and N50 block lengths. Instead we will talk about accuracy and average number of haplotigs per haplotype, two metrics that are more direct representations of the performance of the algorithms and answer two important questions: “How reliable are the results?”, *i.e.* what are the proportions of accurate, erroneous and missing calls? And “How informative are they?”, *i.e.* by how many haplotigs is each haplotype represented? nPhase and WhatsHap polyphase were both applied to our 20X test datasets of different ploidy and heterozygosity levels. nPhase was tested using its default parameters and WhatsHap polyphase was tested with all six possible values of its adjustable sensitivity parameter. We report here the average accuracy, error and missing rates, as well as the average number of haplotigs obtained for the genome, normalized by the ploidy.

In our tests nPhase has an average accuracy of 91.2%, slightly outperforming WhatsHap polyphase’s most sensitive setting (5), which yields an average accuracy of 90.1%, and its second most sensitive setting (4) which yields an average accuracy of 88.9% (Figure 5a). Lower sensitivity levels for WhatsHap polyphase quickly lose accuracy, with the next lowest setting yielding only 81.1% accuracy on average, and its least sensitive setting only reaching 59% accuracy.

In addition to its high accuracy, nPhase is highly contiguous, outputting these accurate results, on average, in 3.4 haplotigs per chromosome per haplotype (Figure 5b). The highly accurate WhatsHap polyphase sensitivity levels (5 and 4) output their results in a highly discontiguous 258.7 and 88.9 haplotigs per haplotype, respectively. In order to output results of similar contiguity to nPhase, WhatsHap polyphase must sacrifice accuracy and drop to a sensitivity level of 1 or 0, which output 2.5 and 0.9 haplotigs per chromosome per haplotype, respectively. This tradeoff between accuracy and contiguity performed by WhatsHap polyphase does not appear to have a useful middle ground and nPhase demonstrates that it is not necessary to make a choice given that it simultaneously achieves both.

## Discussion

We developed nPhase, an algorithm that relies on a few intuitive rules to process an input dataset of long reads, reduced to heterozygous positions, outputting as few clusters as possible which we have shown correspond to the true haplotypes with >90% accuracy. By not specifying the ploidy of the sample in any step, we allow nPhase to adapt to the particularities of the dataset and do not run the risk of forcing an incorrect result to fit such an arbitrary algorithmic constraint. We provide nPhase as part of a pipeline that enables anyone to use their short and long read sequences of the same sample as inputs and obtain a list of SNPs and a fastQ file for each predicted haplotig.

Through our validation tests, we determined that there is a set of parameters for nPhase that performs optimally in nearly all of our test cases and that the algorithm performs well even with very low levels of genetic distance between haplotypes. We found that as little as 10X coverage can yield satisfying results. More complex cases, such as when there is a high ploidy coupled with a low heterozygosity, should benefit from higher coverage and a more stringent parameter for the minimum overlap (0.25 for example). Further investigation would be needed in order to more adequately define how these difficult samples should be treated. We also demonstrated with our benchmarking tests that nPhase outputs far more accurate and contiguous haplotigs than alternative polyploid phasing methods.

As an alignment-based phasing algorithm, the performance of nPhase is going to be highly dependent on the quality of the reference genome being mapped against. Consequently, structural variants between the sample and the reference, or even structural variants within the sample are presently not explicitly identified and phased by the algorithm. In order to resolve structural variants between the sample and the reference or even between haplotypes in the sample, we need to rely on the information in split reads. Here, we used a simple strategy to stitch together some of the haplotigs we obtain without using all of the information contained within split reads. Leveraging the full potential of split reads is a crucial next step to improve the contiguity of phased blocks. The main difficulty in using split reads appears to be that these alignments are significantly less reliable and will need to be processed differently to account for that.

We made the choice not to base our phased blocks on insertion or deletion information. This information can still be obtained in the phased blocks by generating a *de novo* assembly using nPhase’s fastQ output for the relevant haplotig and could be integrated in future developments.

With the nPhase algorithm we believe that the problem of switch errors in polyploid phasing is largely solved, the next important hurdle for polyploid phasing is finding an appropriate way to handle split reads to solve the remaining problems of contiguity and structural variants both within a sample and between the sample and the reference we align to. nPhase can still be used as a preprocessing step for any study of phased polyploid SVs and INDELs since that information is partially held within its output of fastQ files of phased reads.

Overall, nPhase provides, for the first time, an accurate and contiguous picture of polyploid genomes using only a reference genome and short and long reads. It paves the way for a better understanding of the origins of hybrid polyploid organisms, the true diversity of polyploid populations with potential hints on their origins and their relation to other diploid or haploid strains, and provides a clearer picture to investigate phenotypic effects tied to alleles which were previously inaccessible to us.

## Methods

### Total DNA extraction

Single colonies of each natural isolate were isolated by streaking on YPD media, containing ampicillin (50 μg/mL). Cells from one colony of each isolate were grown in 60 mL of YPD at 30°C for 24 hours. We extracted the total DNA of each isolate using the QIAGEN Genomic-tip 100/G kit, according to manufacturer’s instructions.

### Library preparation

We used the EXP-NBD104 native barcoding kit (Oxford Nanopore) and the protocol provided by the manufacturer to barcode the total DNA of each of the isolates. The barcoded DNA was then quantified with a Qubit^®□^ 1.0 fluorometer (Thermo Fisher) and pooled together with an equal amount of DNA coming from each isolate. We then used the SQK-LSK109 ligation sequencing kit (Oxford Nanopore) to finish the library preparation. Finally, the library was loaded to a R9.3 flow cell for a 72 hour run.

### Data pre-processing

The short reads are mapped to a reference genome using bwa^25^ with the command bwa mem -M. We ran GATK^26^ MarkDuplicates then variant called with GATK 4.0’s HaplotypeCaller --ploidy 2 to identify heterozygous positions. Long reads are basecalled, adapter trimmed and demultiplexed by Guppy. They are then mapped to the same reference using NGMLR^27^. We keep only primary alignments and split reads with the samtools^28^ flag 260.

We determine the positions of heterozygous SNPs from the VCF obtained by GATK by looking for positions where AF=1.00 in the file. We reduced each long read to the set of variable positions it overlaps (Supplemental Figure 2b). To simplify later computational steps, we remove long reads that are subsets of other long reads.

nPhase is only capable of phasing SNPs if they are identified by the variant calling step. This is not necessarily always the case, and the accuracy metrics are based on the total number of SNPs identified in the polyploid sample by the variant calling step. However, unidentified SNPs will still exist in the reads, so if the algorithm performs a proper clustering of the reads the information will still be available and readily extracted by a closer view of the results.

### Context coverage

Long reads are error-prone but it is important not to perform any form of error correction to ensure that the heterozygosity is not incorrectly flattened or mis-assigned. The nPhase pipeline works with raw long reads. In order to minimize the influence of these errors we consider that SNP coverage is a useful indicator of quality. We count the number of times each heterozygous SNP is present in a specific context in our dataset. We define context as being the closest flanking heterozygous SNPs (two heterozygous SNPs upstream and two heterozygous SNPs downstream). The context information will be used to better inform the nPhase algorithm and allow it to escape the situation where a sequencing error randomly converts a well-supported SNP to another SNP that is well-supported in another haplotype (Supplemental Figure 2b).

### Output results

Once nPhase is done running it outputs several files:

i. A fastQ file for each haplotig containing all of the reads that have been clustered together for this haplotig, this file can then be used with a *de novo* assembly or alignment tool for further analysis.
ii. A tab separated file listing the consensus base for each heterozygous position contained within each haplotig. There are three columns: chromosome, position and consensus base. If two different bases are equally represented for a given position and equally well supported within the cluster they will both be represented in this file on separate lines. This file is sorted by position.
iii. A plot representing the different haplotigs along the reference genome, similar to the one displayed in Figure 3 but lacking the haplotype color code since the ground truth is not known in a typical use case of nPhase.

### nPhase parameter description

nPhase has a total of 4 parameters which can be adjusted to better fit the sample. These parameters are the following (Supplemental Figure 3):

#### S, the minimum fraction of similarity between two reads

When two reads overlap with each other we calculate their similarity by dividing the number of heterozygous SNPs they share by the number of heterozygous positions they both cover. If that fraction is smaller than the parameter S, then we will consider that these two reads cannot be part of the same haplotype. This parameter can be set to any fraction between 0 and 1, by default it is set at 0.01, or 1% similarity.

#### O, the minimum fraction of overlap between two reads

When two reads overlap with each other we can count the number of heterozygous positions they both cover. If they both cover more than 100 heterozygous positions, this parameter is ignored. If they cover fewer than 100 heterozygous positions then we calculate the overlap by dividing the number of heterozygous positions the two reads have in common by the total number of heterozygous positions covered by the smaller of the two reads. In this case, smaller does not necessarily mean a shorter read, it means a read that covers fewer heterozygous positions. If this overlap is smaller than the parameter O, then we consider that these two reads do not overlap enough for us to conclusively determine if they’re part of the same haplotype. This parameter can be set to any fraction between 0 and 1, by default it is set at 0.1, or 10% overlap.

#### L, the minimum number of reads supporting a haplotig

Once nPhase has clustered all of the reads into different haplotigs, the user may want to filter out all haplotigs that are supported by fewer than N reads.This parameter can be set to any integer N ≥ 0, by default it is set at 0. If set to N, it will not output any cluster supported by fewer than N reads.

#### ID, the maximum amount of change when merging clusters

When nPhase considers merging two clusters of reads into one new cluster it must determine if these two clusters are similar enough to warrant merging them together or if they should remain unique clusters, representative of unique haplotypes. Since these are clusters, every heterozygous position is potentially covered multiple times, sometimes with different reads in the same cluster indicating conflicting bases for the same position. We can calculate the number of reads voting for each base in a given cluster and determine the “demographics” for that position. We can take this further and have an overview of every heterozygous position in the cluster and how well-supported each base is. The base that has the majority of support is considered to be the “true” base for that cluster. When we merge two clusters together, we potentially change these “demographics”. These changes either further strengthen the position of the majority base for a given position, in which case there is no negative change in the cluster’s “identity” or they weaken the majority base’s position and cause a negative change to the cluster’s “identity”. When there are negative changes to the cluster’s “identity” we can calculate the amount of change that has occurred and if that amount is too high the clusters are not allowed to merge. This parameter can be set to any fraction between 0 and 1, by default it is set at 0.05, or a 5% ID change tolerance.

These parameters are set by default, though they can be modified if needed. The nPhase algorithm will use these parameters as limitations to determine which reads it is allowed to cluster together into haplotigs and which clusters of reads it can merge together into longer haplotigs. Ideally, only the ID parameter needs to be modified, keeping all other parameters very low and forcing the algorithm to merge clusters as aggressively as allowed by the ID parameter.

### Identifying optimal parameters

In order to determine which parameters nPhase should use by default, it is important to understand how these parameters affect the results. Ideally, we will find that there is a set of parameters which is optimal for all possible combinations of ploidy and heterozygosity level, such a set would then become the default recommended parameters for nPhase. If no such set of parameters appears to exist, the next best case is to minimize the impact of as many of the available parameters as possible in order to reduce the parameter a user would need to explore when using nPhase to phase their dataset.

Through our tests, we find that there is a narrow range in the parameter space that results in the optimal performance of nPhase. Intuitively, the optimal strategy appears to be to set the minimum similarity and minimum overlap parameters down to a low value so that all of the reads in the dataset are allowed to merge into a cluster, and to only worry about finding an appropriate threshold for the ID change parameter. Since the ID change parameter controls how dissimilar two clusters need to be in order to be considered two different haplotypes, it is fitting for this parameter alone to have the most pronounced impact on the quality of results. If set too low nPhase will consider small sequencing errors to be evidence of alternate haplotypes, and if set too high it will allow different haplotypes to merge into chimeric and wrong results.

To demonstrate this, we ran nPhase 125 times on 24 different samples of varying coverage, ploidy and number of heterozygous SNPs for a total of 3000 tests. These 125 tests represent every possible combination of the minimum similarity S, minimum overlap O, and maximum identity change ID parameters for the following values: 0.01, 0.05, 0.1, 0.15, 0.25.

The L parameter was set to 0 for these tests since it’s intended for use to clean up results by removing small, lowly supported haplotigs and we wanted to determine how nPhase performs without throwing away any of the data.

We found that S, the minimum similarity parameter, had no influence on the results at these levels (Supplemental Figure 4a). O, the minimum overlap parameter, needs to be at least at 0.1 and seems to show very minor improvements in accuracy at higher levels (Supplemental Figure 4b). The ID parameter has the most influence on the accuracy of the results, with values of 0.05 and 0.1 yielding the best results (Supplemental Figure 4c).

We then looked at the effects of O and ID on the average number of haplotigs per chromosome per parent. We found that the number of haplotigs slightly increases with O (Supplemental Figure 5a), while ID has a strong effect on the contiguity of the results (Supplemental Figure 5b). A higher value for ID leads to a more contiguous assembly, though this comes at the cost of accuracy (Supplemental Figure 5c). We again find that values held between 0.05 and 0.1 provide good results. If we separate our tests by ploidy we can see that, as the ploidy increases the optimal choice for the ID parameter narrows down around 0.05 (Supplemental Figure 6).

Based on our tests, we find that the following set of parameters is the best adapted to handle any sample: S=0.01, O=0.1, L=0, ID=0.05. We use these as our default parameters.

### Influence of coverage

We sought to establish the effects of coverage on the quality metrics of nPhase’s predictions. To do so we performed our tests on a 10X per haplotype dataset and a 20X per haplotype dataset. We found that both accuracy and contiguity are improved by the higher coverage level of 20X per haplotype (Supplemental Figure 7). This effect is observed across ploidy and heterozygosity levels, though the accuracy effects are more pronounced for higher ploidy, lower heterozygosity level samples.

A low number of haplotigs per haplotype is not always a good sign of high contiguity as it can be compatible with a high rate of chimeric haplotigs. Therefore, we looked at the contiguity effects of coverage for our tests using default parameters, which we have previously determined output accurate results. Based on these tests we were able to confirm that the 20X dataset is more contiguous than the 10X dataset (Supplemental Figure 7b). We therefore used the 20X datasets as part of our default analysis.

### Split read stitching step

Some reads align to two or more very distant sequences in the reference genome. These reads can represent a structural variation between the sample and the reference being mapped to. We split them into the different segments that align to the reference and refer to them to as split reads.

Split reads can be very misleading and trusting them blindly would result in chimeric haplotigs. We developed a simple pre-processing strategy to integrate part of the information contained by these split reads.

We run nPhase a first time to obtain our initial haplotigs. We expect some of the edges of haplotigs to correspond to structural variants such as inversions or large INDELs so we identify the SNPs at the edges of these clusters. These are the SNPs which we expect to be included in the split reads that can connect two haplotigs separated by a structural variant, so they are the most trustworthy SNPs in our split read dataset. We reduce each split read to only the heterozygous SNPs that overlap with these regions and re-run the nPhase algorithm with these reads included. Clusters are currently not allowed to combine reads from different reference chromosomes, so split reads can only be used to improve the contiguity of haplotigs on the same reference chromosome.

As described, nPhase does not exploit the information contained in split reads to the fullest extent, only attempting to improve contiguity by stitching together haplotigs on the same chromosome. Once there are only a few remaining haplotigs, further improving contiguity necessarily means stitching longer haplotigs together. This presents a very real danger of creating chimeric haplotigs that have very strong negative effects on accuracy. To validate the usefulness of these steps and this method of using the split read data we ran 3000 tests of nPhase both with split read information and 3000 tests without in order to determine the effects of our split read stitching strategy on both contiguity and accuracy. We found that the contiguity did significantly improve across all of our tests that included split read information, compared to those that did not (Supplemental figure 8a). Encouragingly, when comparing the accuracy distributions of the two sets of tests they are virtually identical (Supplemental figure 8b). The tests that used split reads were very slightly less accurate than their counterparts but much more contiguous, motivating our decision to integrate the use of split read information in nPhase.

### Performance limits

With default parameters the nPhase algorithm took between 1 minute and nearly 5 hours of runtime on a single CPU (the nPhase algorithm has not been parallelized), and between 0.6 GB and 31.8 GB of memory (Supplemental table 3). The runtime and memory usage are clearly tied to the ploidy and heterozygosity level. A higher ploidy and higher heterozygosity level translates to a significant increase in runtime and memory usage. Each diploid test, up to 1% heterozygosity, ran in less than an hour and ten minutes and used less than 8 GB of memory. Triploid tests took a minimum of 3.5 minutes of CPU time and 0.9 GB of memory to run for the 0.05% heterozygosity level example, and a maximum of three hours and ten minutes of CPU time and 19 GB of memory for the 1% heterozygosity level test. The tetraploid examples were the most resource intensive, using up a minimum of 6 minutes of CPU time and 1.25 GB of memory for the 0.05% heterozygosity level, and a maximum of four hours and fifty minutes of CPU time and 31.8 GB of memory to run. nPhase can output results in a reasonable time using moderate memory resources. If run on a particularly large genome in a time-sensitive context, nPhase could be applied to individual chromosomes in parallel. It’s also reasonable to consider down-sampling the number of SNPs to a heterozygosity level of around 0.5% given the results obtained are comparable and run in less than half the time as the 1% heterozygosity level tests. All of the heterozygous SNPs would still be present in the long reads and could be recovered from the fastQ files associated to the predicted haplotypes.

## Data Availability

The nPhase algorithm and the nPhase pipeline are both available at https://github.com/nPhasePipeline/nPhase

Oxford Nanopore sequencing data is available under the study accession number <u>PRJEB39456</u>

Illumina short read data is taken from the 1,011 yeast genomes project and their SRA accession numbers are given in Supplemental Table S1.

## Acknowledgments

This work was supported by a National Institutes of Health (NIH) grant R01 (GM101091-01), an Agence Nationale de la Recherche (GM101091-01) grant and a European Research Council (ERC) Consolidator grant (772505).

## References

1. Mahmoud, M. et al. Structural variant calling: the long and the short of it. Genome Biol 20, (2019).

2. Sohn, J. & Nam, J.-W. The present and future of de novo whole-genome assembly. Brief Bioinform 19, 23–40 (2018).

3. Kitzman, J. O. et al. Haplotype-resolved genome sequencing of a Gujarati Indian individual. Nat. Biotechnol. 29, 59–63 (2011).

4. Roach, M. J. et al. Population sequencing reveals clonal diversity and ancestral inbreeding in the grapevine cultivar Chardonnay. PLoS Genet. 14, e1007807 (2018).

5. Hamazaki, K. & Iwata, H. Haplotype-based genome wide association study using a novel SNP-set method□: RAINBOW. bioRxiv 612028 (2019).

6. Sanjak, J. S., Long, A. D. & Thornton, K. R. A Model of Compound Heterozygous, Loss-of-Function Alleles Is Broadly Consistent with Observations from Complex-Disease GWAS Datasets. PLOS Genetics 13, e1006573 (2017).

7. Benitez, J. A., Cheng, S. & Deng, Q. Revealing allele-specific gene expression by single-cell transcriptomics. Int. J. Biochem. Cell Biol. 90, 155–160 (2017).

8. Wagner, N. D., He, L. & Hörandl, E. Relationships and genome evolution of polyploid Salix species revealed by RAD sequencing data. bioRxiv 864504 (2019).

9. Eriksson, J. S. et al. Allele phasing is critical to revealing a shared allopolyploid origin of Medicago arborea and M. strasseri (Fabaceae). BMC Evol Biol 18, 9 (2018).

10. Yang, J. et al. Incomplete dominance of deleterious alleles contributes substantially to trait variation and heterosis in maize. PLOS Genetics 13, e1007019 (2017).

11. Fay, J. C. et al. A polyploid admixed origin of beer yeasts derived from European and Asian wine populations. PLOS Biology 17, e3000147 (2019).

12. Zhou, R.-N. & Hu, Z.-M. The Development of Chromosome Microdissection and Microcloning Technique and its Applications in Genomic Research. Curr Genomics 8, 67–72 (2007).

13. Snyder, M. W., Adey, A., Kitzman, J. O. & Shendure, J. Haplotype-resolved genome sequencing: experimental methods and applications. Nature Reviews Genetics 16, 344–358 (2015).

14. Zhang, X., Wu, R., Wang, Y., Yu, J. & Tang, H. Unzipping haplotypes in diploid and polyploid genomes. Comput Struct Biotechnol J 18, 66–72 (2019).

15. Koren, S. et al. De novo assembly of haplotype-resolved genomes with trio binning. Nat Biotechnol (2018).

16. He, D., Saha, S., Finkers, R. & Parida, L. Efficient algorithms for polyploid haplotype phasing. BMC Genomics 19, 110 (2018).

17. Browning, S. R. & Browning, B. L. Rapid and Accurate Haplotype Phasing and Missing-Data Inference for Whole-Genome Association Studies By Use of Localized Haplotype Clustering. The American Journal of Human Genetics 81, 1084–1097 (2007).

18. Patterson, M. et al. WhatsHap: Weighted Haplotype Assembly for Future-Generation Sequencing Reads. Journal of Computational Biology 22, 498–509 (2015).

19. Chin, C.-S. et al. Phased diploid genome assembly with single-molecule real-time sequencing. Nat. Methods 13, 1050–1054 (2016).

20. Schrinner, S. D. et al. Haplotype Threading: Accurate Polyploid Phasing from Long Reads. bioRxiv 2020.02.04.933523 (2020).

21. Xie, M., Wu, Q., Wang, J. & Jiang, T. H-PoP and H-PoPG: heuristic partitioning algorithms for single individual haplotyping of polyploids. Bioinformatics 32, 3735–3744 (2016).

22. Motazedi, E., Finkers, R., Maliepaard, C. & de Ridder, D. Exploiting next-generation sequencing to solve the haplotyping puzzle in polyploids: a simulation study. Brief. Bioinformatics 19, 387–403 (2018).

23. Moeinzadeh, M.-H. et al. Ranbow: A fast and accurate method for polyploid haplotype reconstruction. PLOS Computational Biology 16, e1007843 (2020).

24. Peter, J. et al. Genome evolution across 1,011 *Saccharomyces cerevisiae* isolates. Nature 556, 339–344 (2018).

25. Li, H. & Durbin, R. Fast and accurate short read alignment with Burrows–Wheeler transform. Bioinformatics 25, 1754–1760 (2009).

26. Van der Auwera, G. A. et al. From FastQ data to high confidence variant calls: the Genome Analysis Toolkit best practices pipeline. Curr Protoc Bioinformatics 43, 11.10.1–11.10.33 (2013).

27. Sedlazeck, F. J. et al. Accurate detection of complex structural variations using single-molecule sequencing. Nature Methods 15, 461–468 (2018).

28. Li, H. et al. The Sequence Alignment/Map format and SAMtools. Bioinformatics 25, 2078–2079 (2009).

