## Supplemental Figures for "nPhase: An accurate and contiguous phasing method for polyploids"

**Supplemental Figure 1.** Graphical representations of nPhase output results for every genome analyzed.

2n dataset, 0.05% heterozygosity level

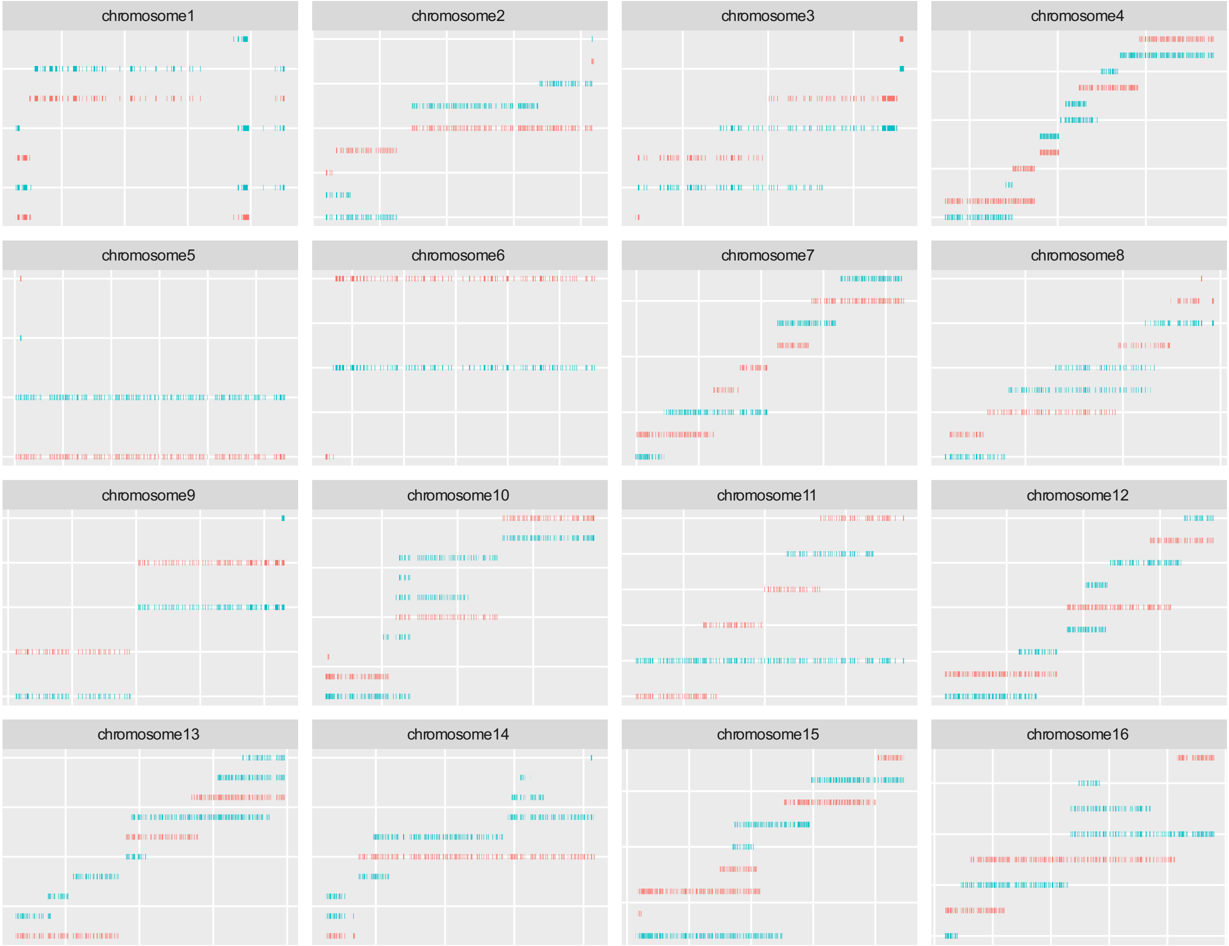

2n dataset, 0.1% heterozygosity level

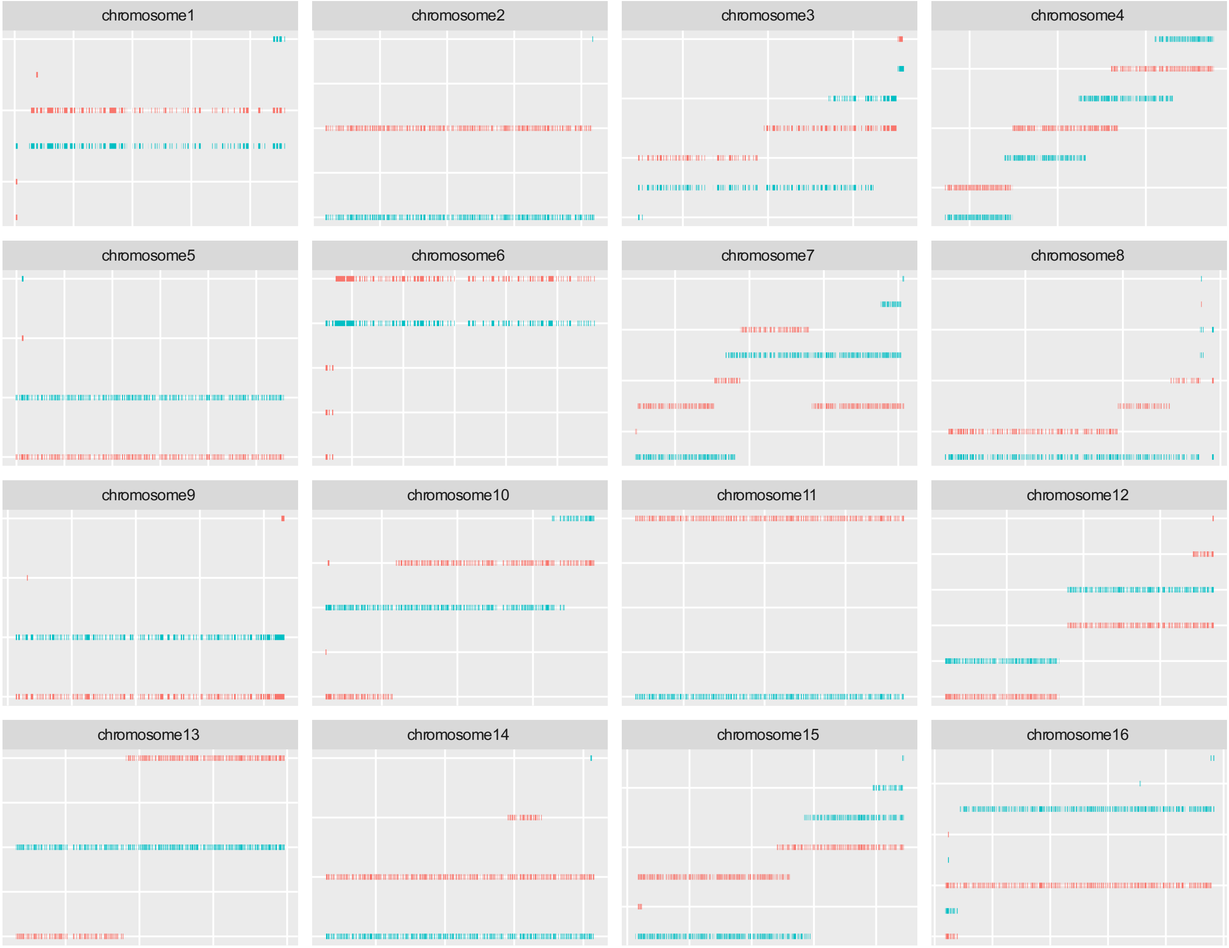

2n dataset, 0.5% heterozygosity level

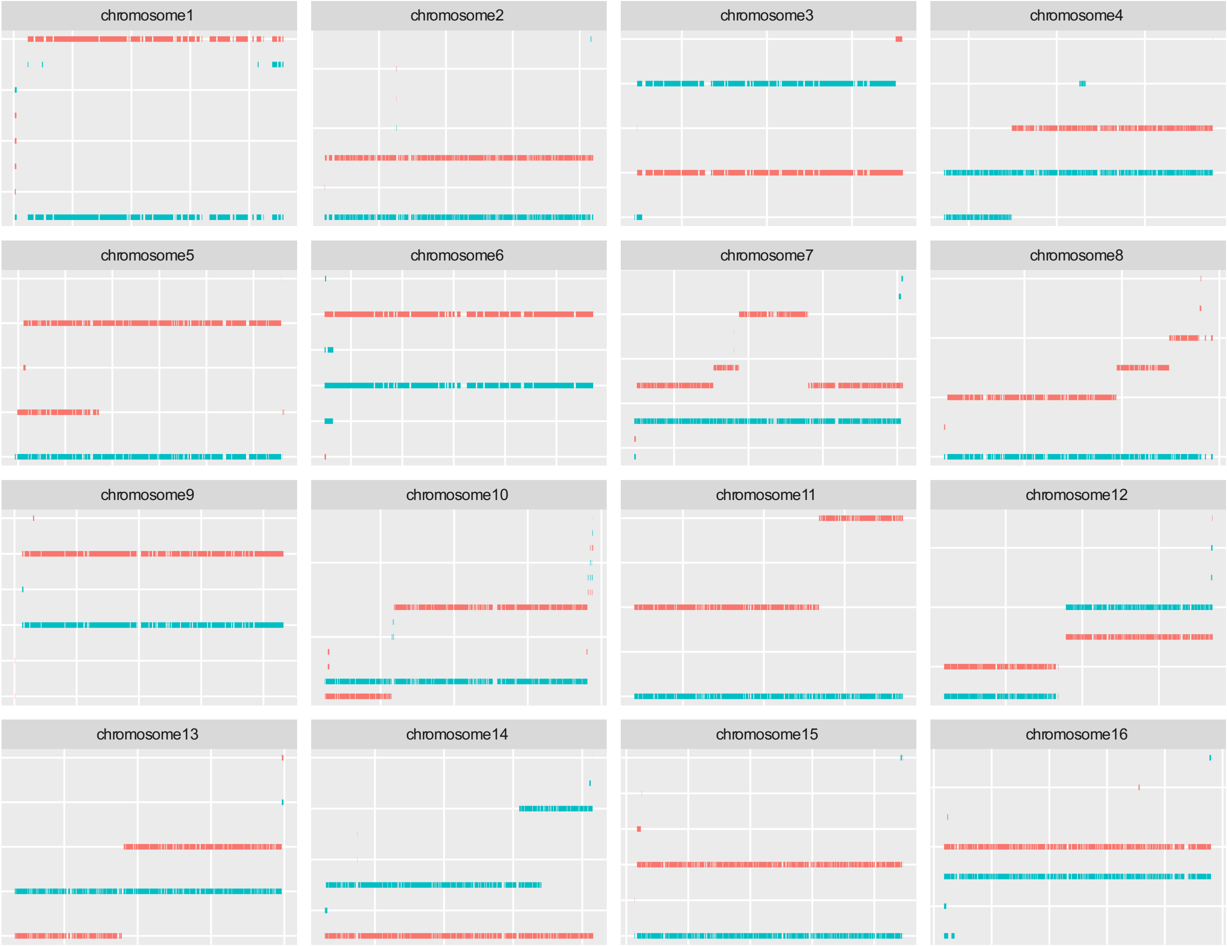

2n dataset, 1% heterozygosity level

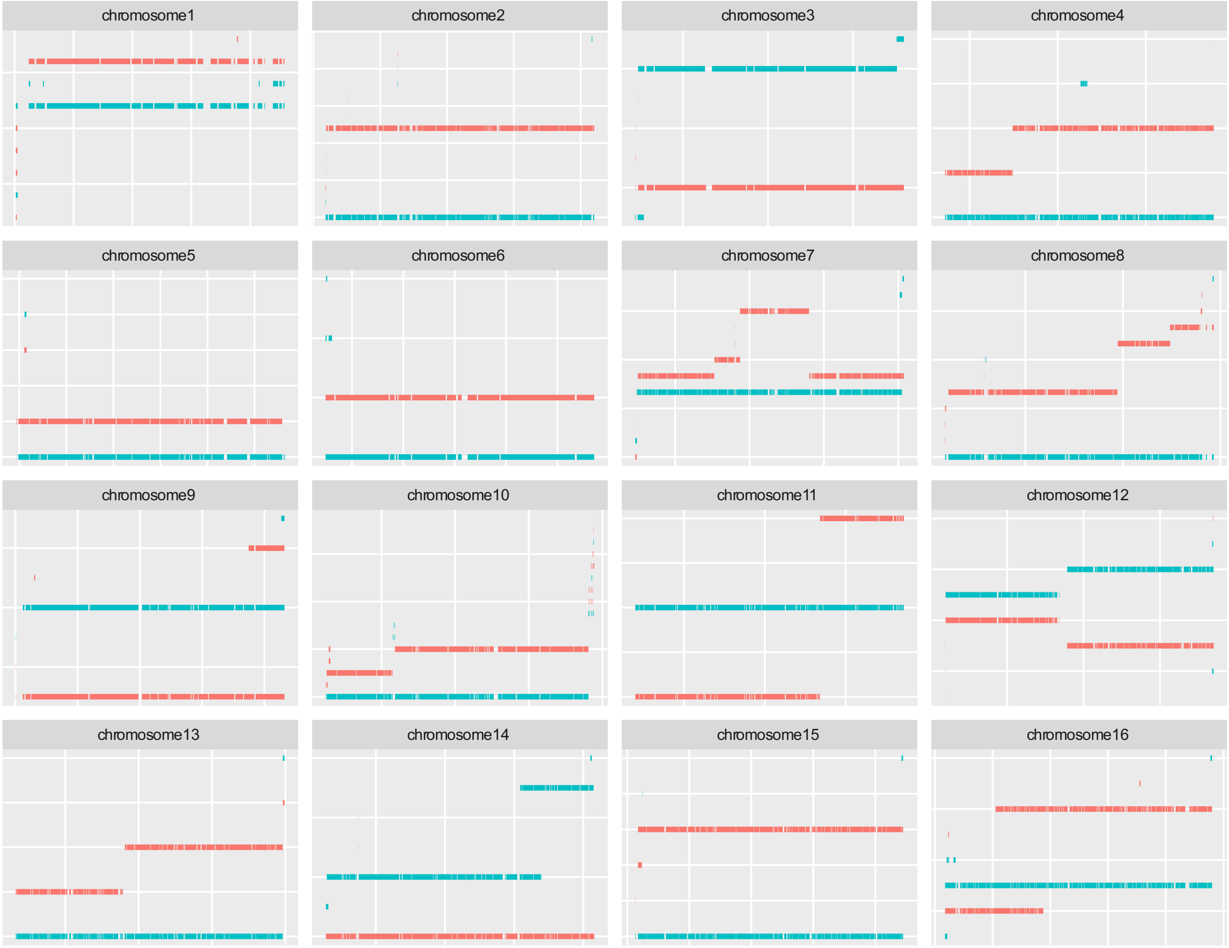

3n dataset, 0.05% heterozygosity level

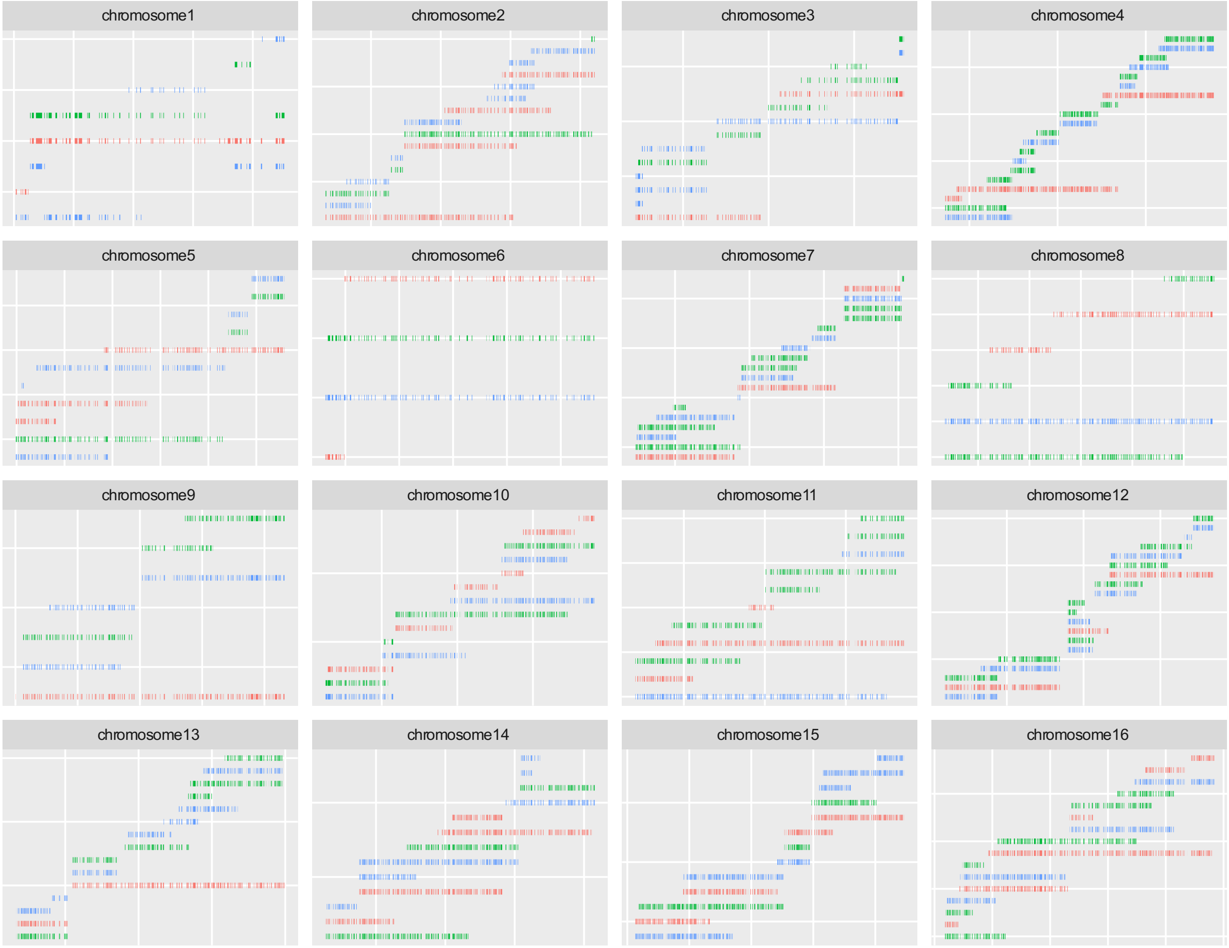

3n dataset, 0.1% heterozygosity level

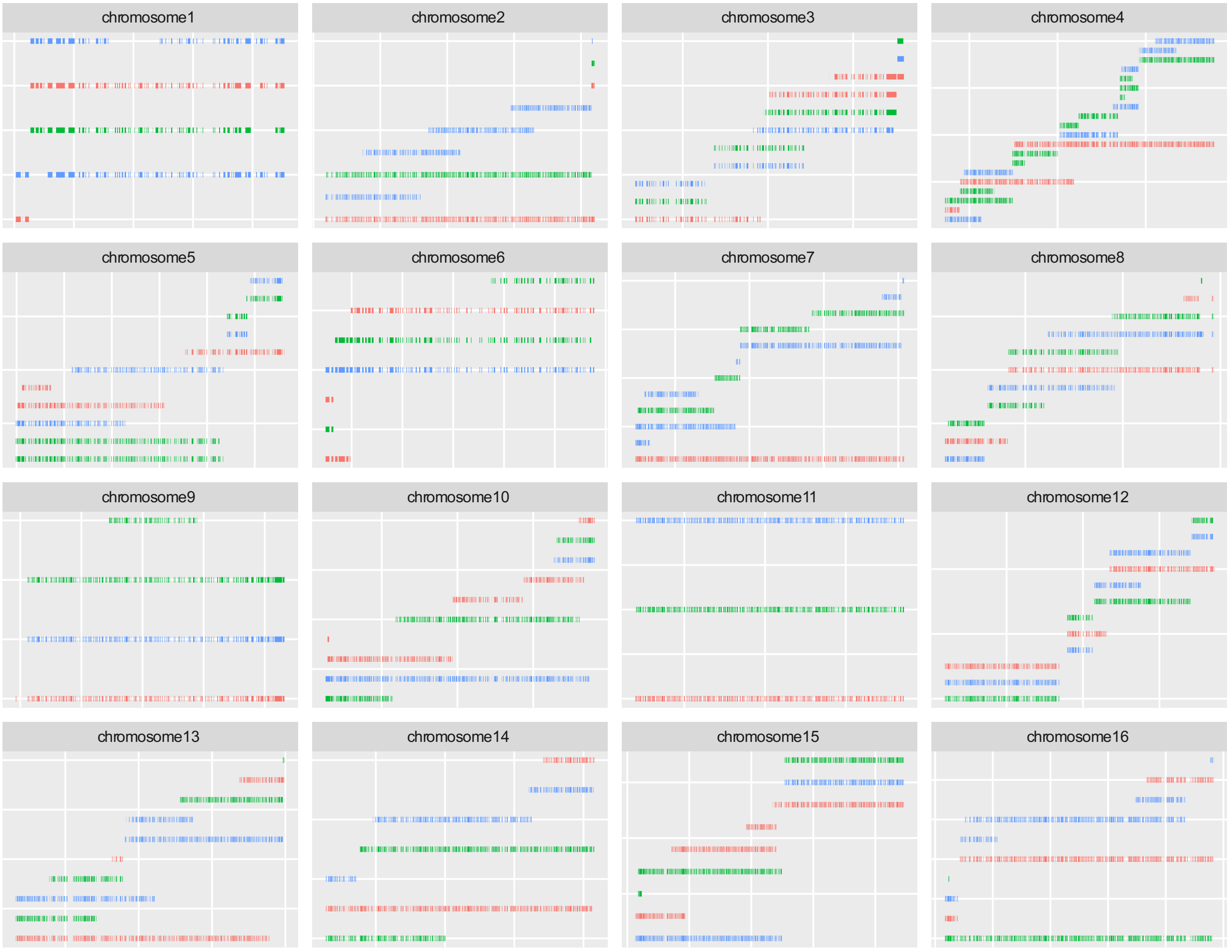

3n dataset, 0.5% heterozygosity level

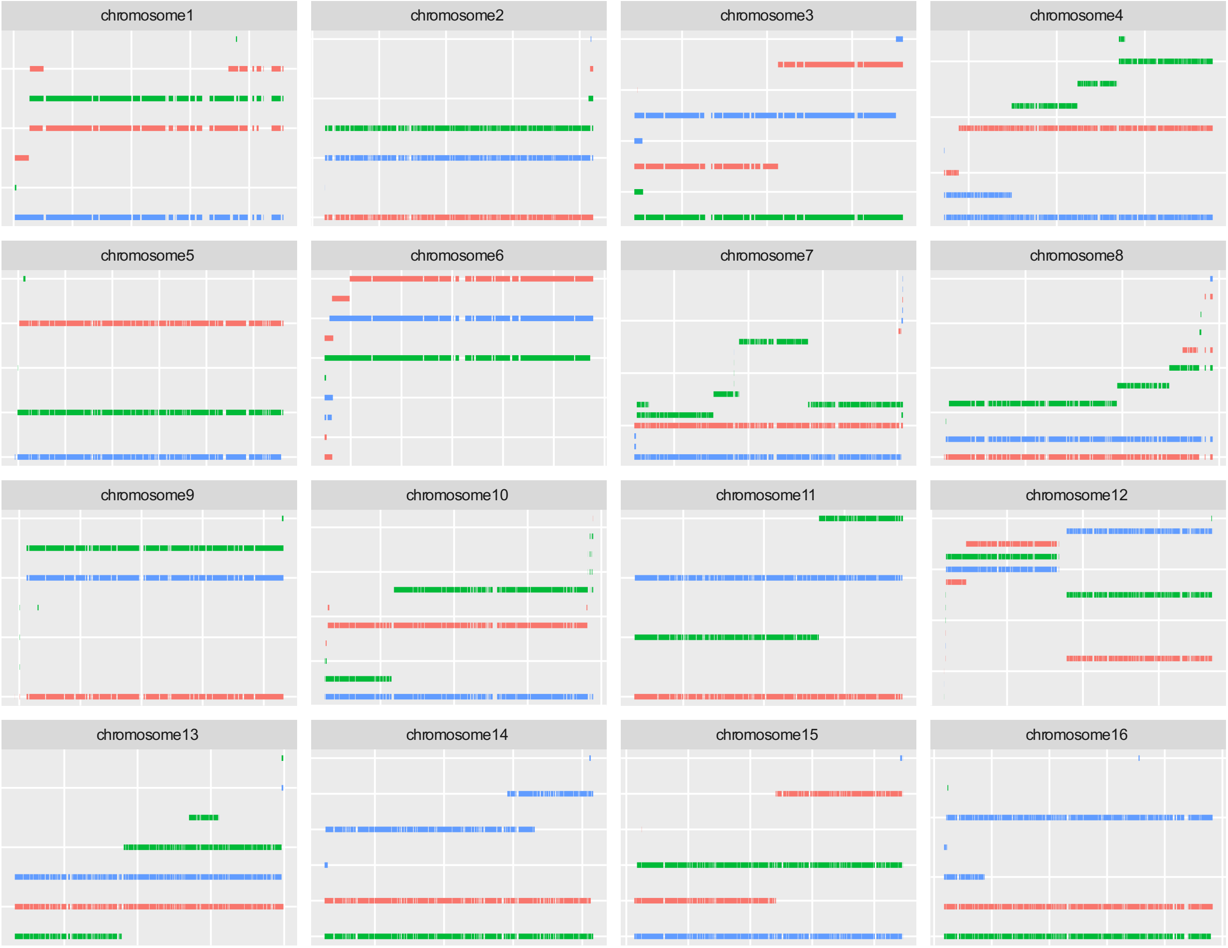

3n dataset, 1% heterozygosity level

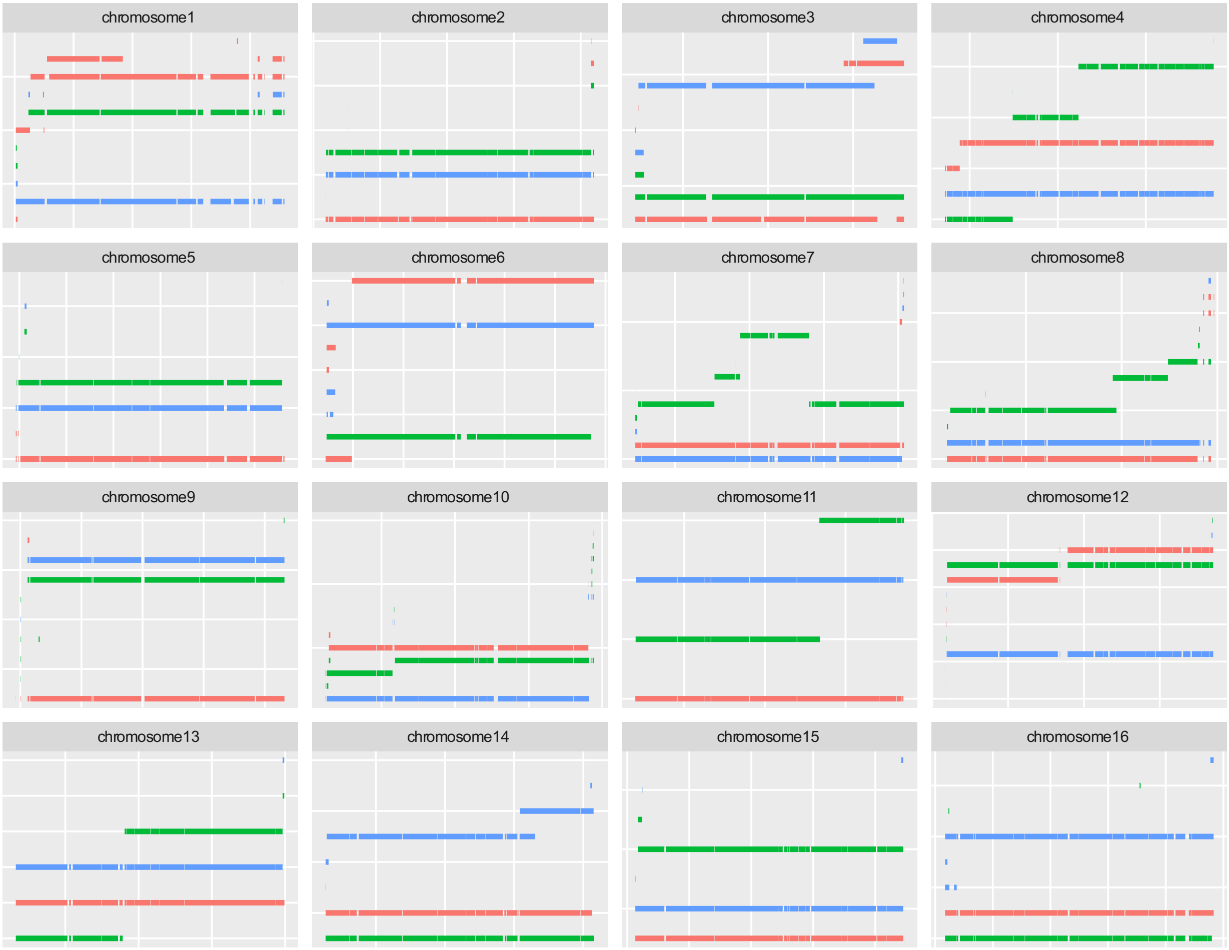

4n dataset, 0.05% heterozygosity level

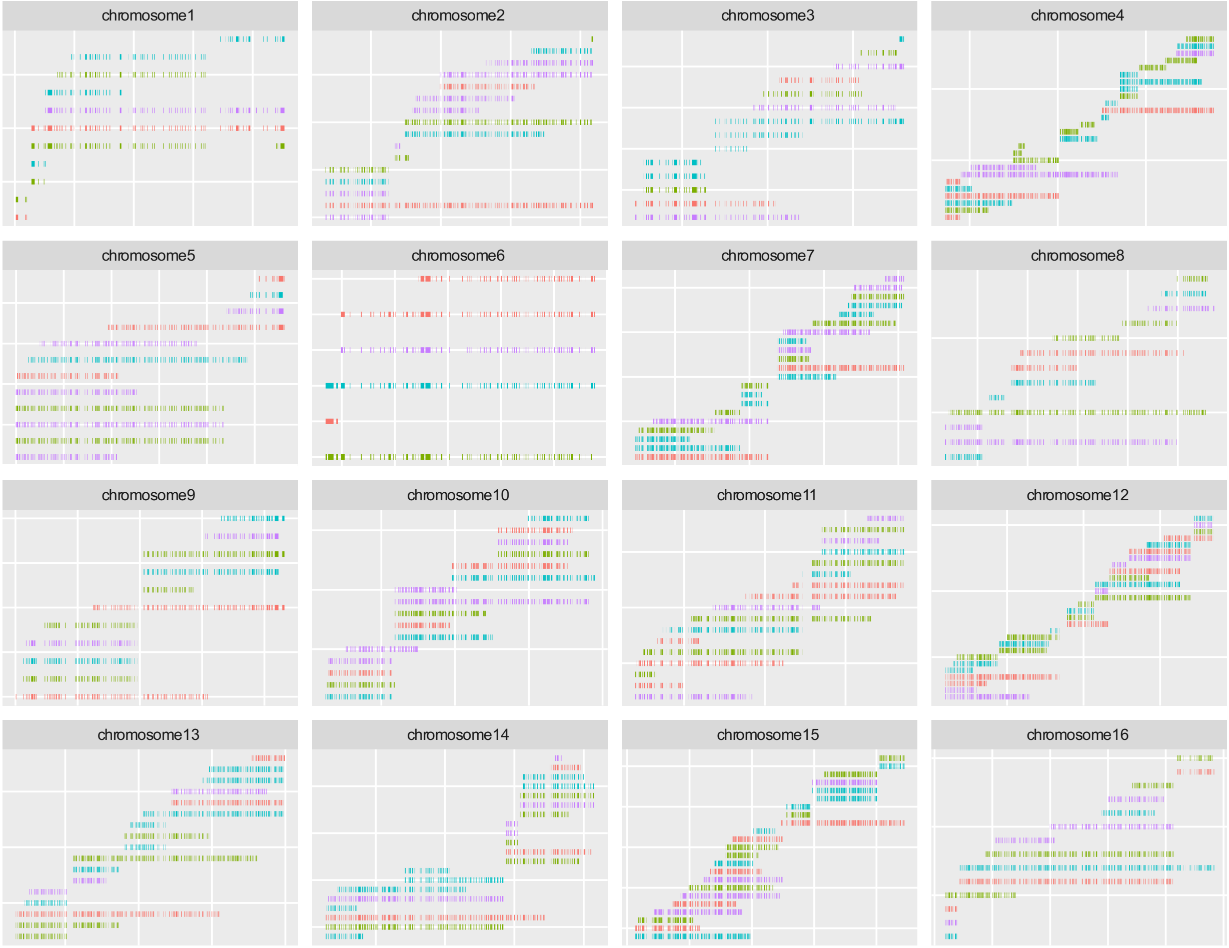

4n dataset, 0.1% heterozygosity level

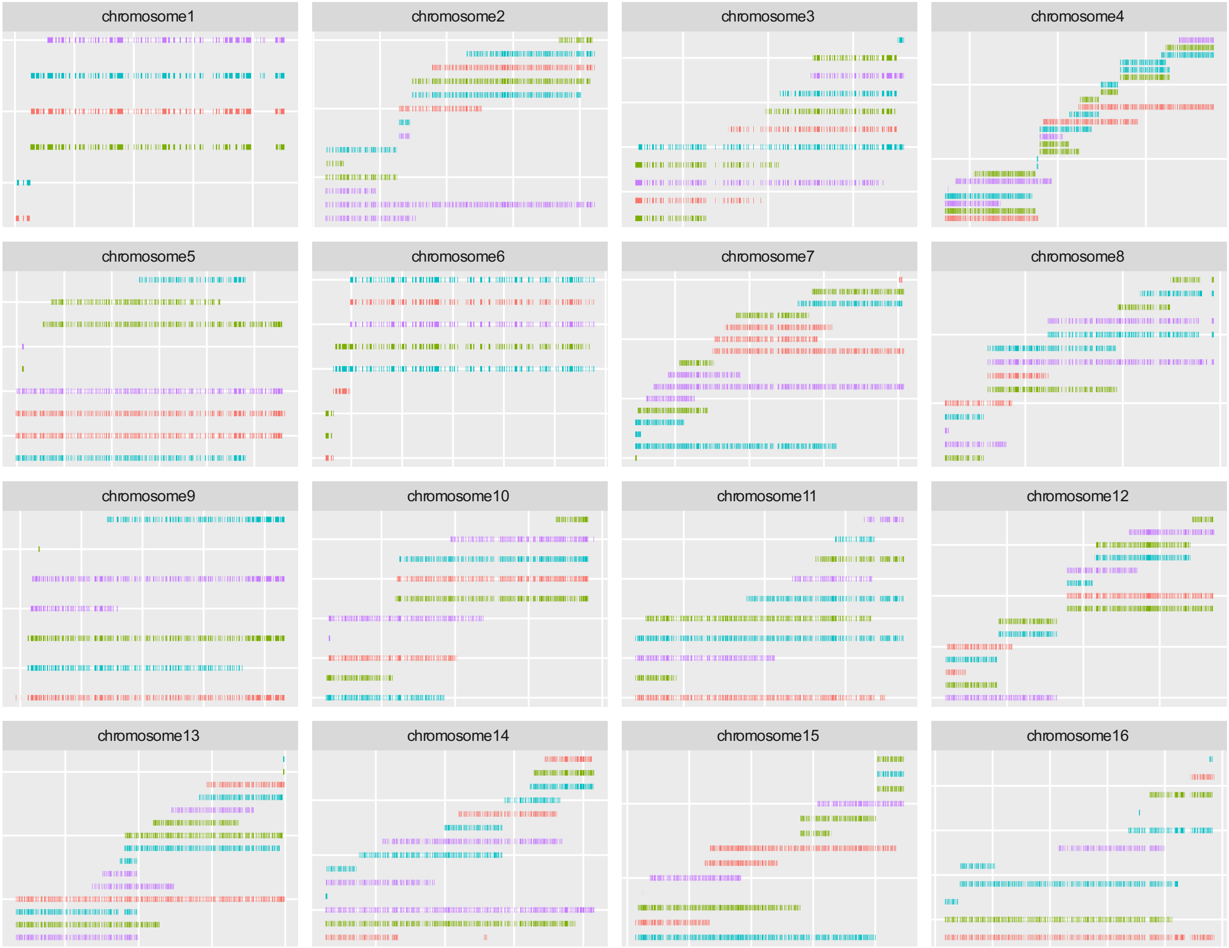

4n dataset, 0.5% heterozygosity level

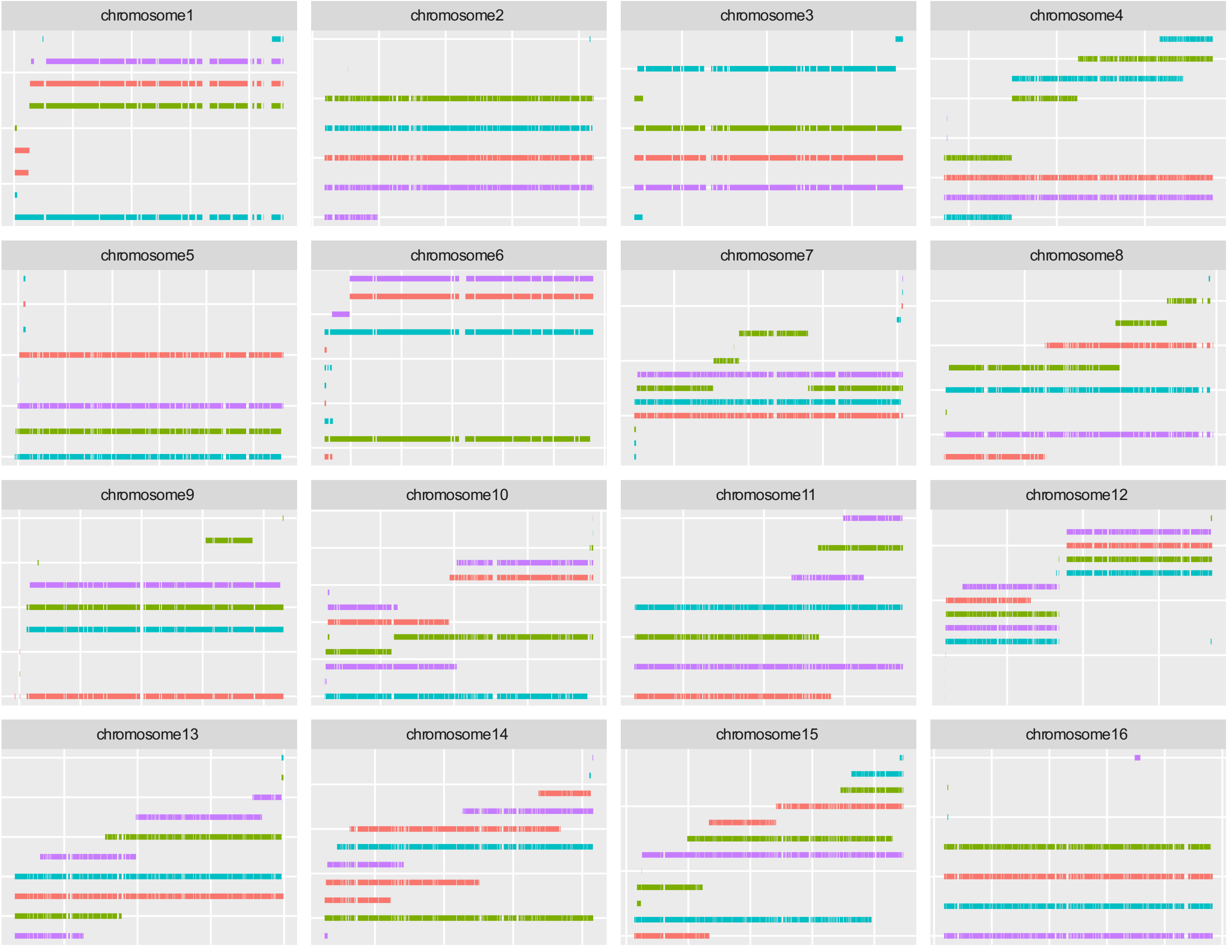

4n dataset, 1% heterozygosity level

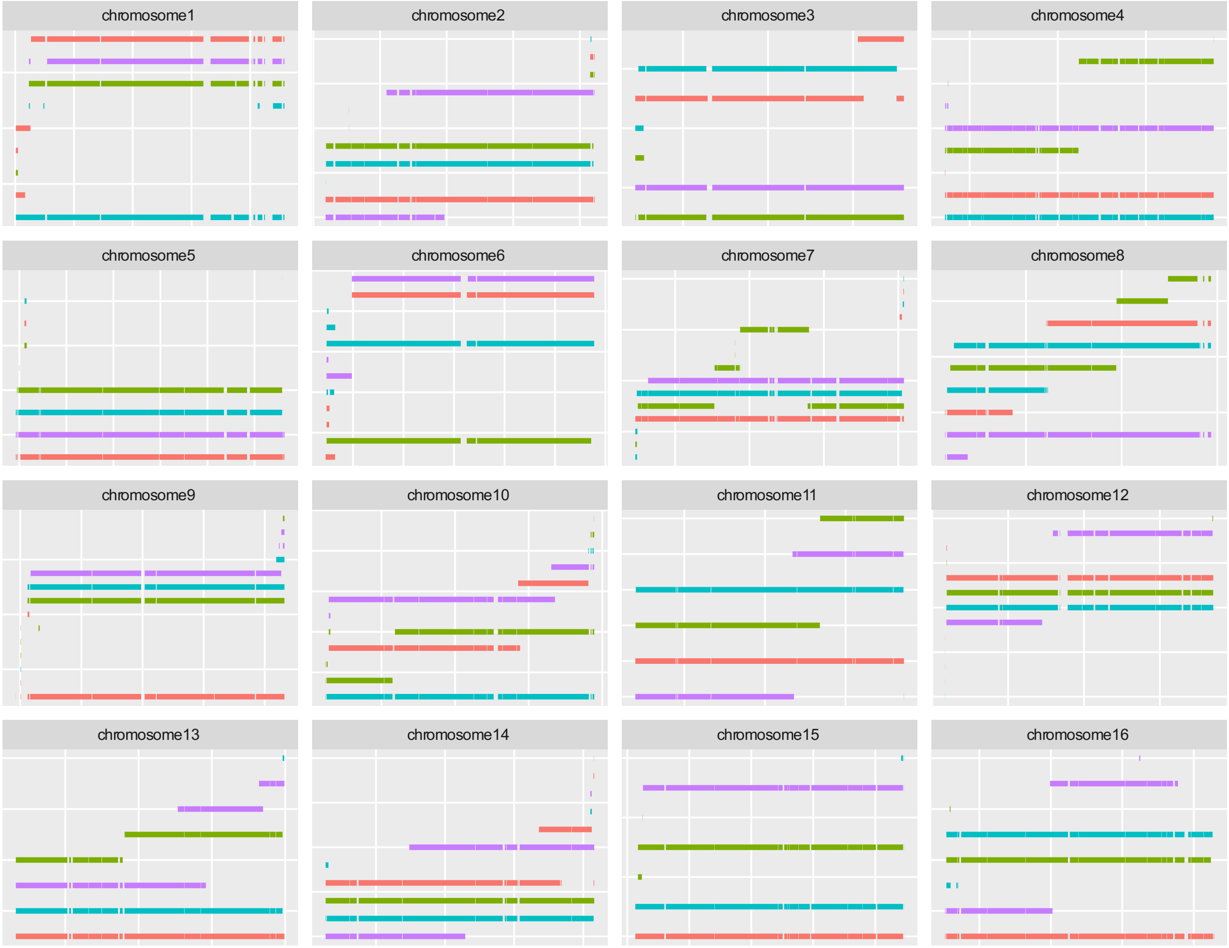

**Supplemental Figure 2.** Long read pre-processing steps**.** (**a**) Simplifying long reads**.** Each long read is reduced to the set of variable positions it overlaps. Hence the first sequence becomes ATC, the second becomes CGA and the third becomes AGC. We keep track of the position and chromosome on which each SNP is found. (**b**) Context coverage**. T** and **G** are equally covered without context, but with context we see that **AGC** and **CTA** are not as highly covered as **ATC** and **CGA**.

(a) (b)

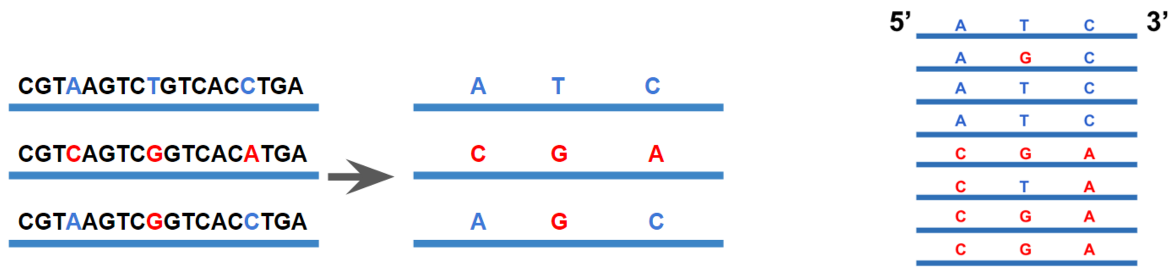


**Supplemental Figure 3.** nPhase parameters. The parameters S, O, L and ID are the only parameters that can be user-set in the nPhase algorithm.

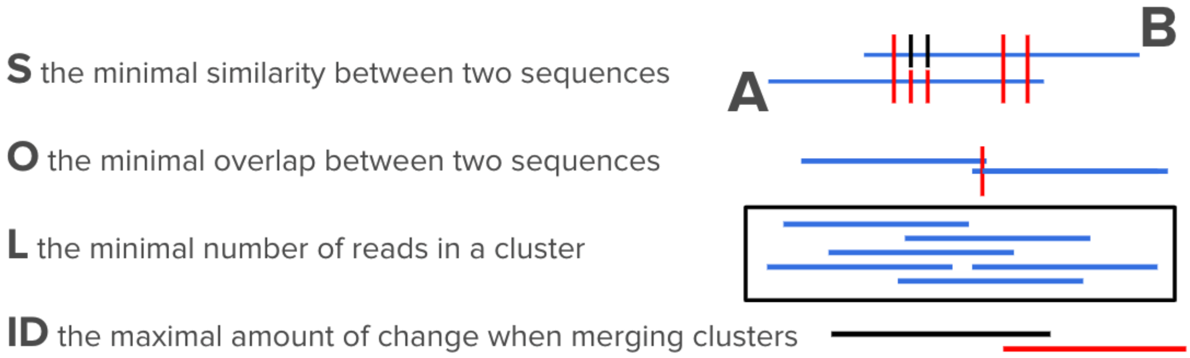

**Supplemental figure 4.** Effects of parameters on prediction accuracy. We ran a total of 3000 tests using different nPhase parameters in order to evaluate their effects on the accuracy of the results. We found that the minimum overlap and minimum similarity parameters had minimal effects as shown by these violin plots of the accuracy for different values of each parameter, whereas the maximum ID parameter was much more influential. (**a**) The violin plots display an optimal performance for minimum overlap values of at least 0.1 , which corresponds to the presence of at least 10% of heterozygous SNPs in common between two clusters. This parameter only has an effect concerning clusters that have fewer than 100 heterozygous SNPs in common. (**b**) The violin plots for the different possible values attributed to the minimum similarity parameter are all the same, suggesting that at these values the parameter has no effect. (**c**) Based on these violin plots we found that, overall, the most reliable value for this parameter is 0.05, i.e. two clusters can only merge if it does not change their demographics by more than 5%. A value of 0.01 led to overall worse results, and higher values seem to split into two groups, with one that maintains a high accuracy and another that further falls as the ID parameter is set to higher and more lenient values.

(a)

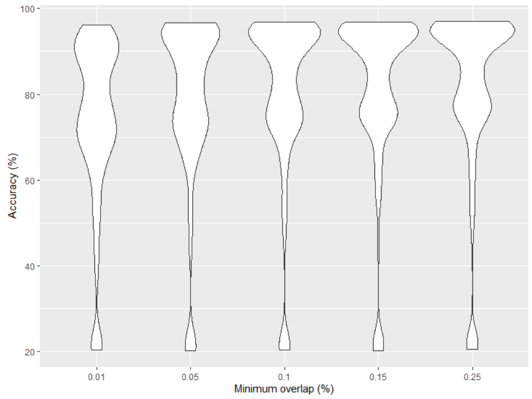


(b)

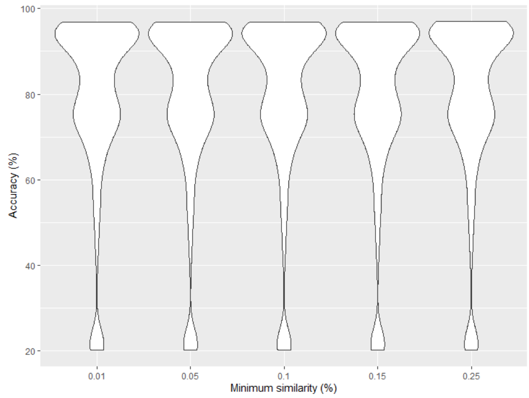


(c)

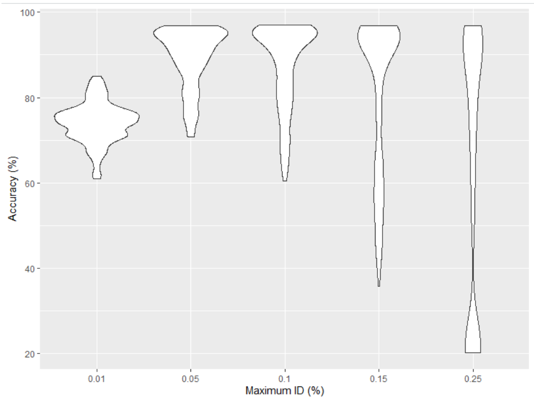

**Supplemental Figure 5.** Effects of parameters on contiguity. We ran a total of 3000 tests using different nPhase parameters in order to evaluate their effects on the contiguity of the results. We found that the minimum had a small effect, whereas the maximum ID parameter was much more influential. (**a**) These violin plots show how different values for the minimum overlap parameter affect the number of haplotigs. The Y axis displays the number of haplotigs per chromosome normalized by the number of haplotypes. We see a weak but predictable increase in the number of haplotigs as we increase this value and make it more stringent, though all parameter values shown here result in very comparable distributions. (**b**) These violin plots show how different values for the maximum ID parameter affect the number of haplotigs. The Y axis displays the number of haplotigs per chromosome normalized by the number of haplotypes. We observe here that the 0.01 value for this parameter, previously shown to lead to inaccurate results, also displays a significantly higher number of haplotigs than other values tested. As we increase the value of the ID parameter, rendering it less stringent, we also lower the number of haplotigs obtained. (**c**) This graph is similar to the one shown in (a), showing the normalized number of haplotigs per chromosome on the Y axis and the different values for the ID parameter in the X axis. We also color coded the individual tests, a lighter color denotes a more accurate result, whereas a darker color denotes a less accurate result. As the maximum ID parameter increases and becomes more lenient, we see that the most contiguous results are significantly less accurate.

(a)

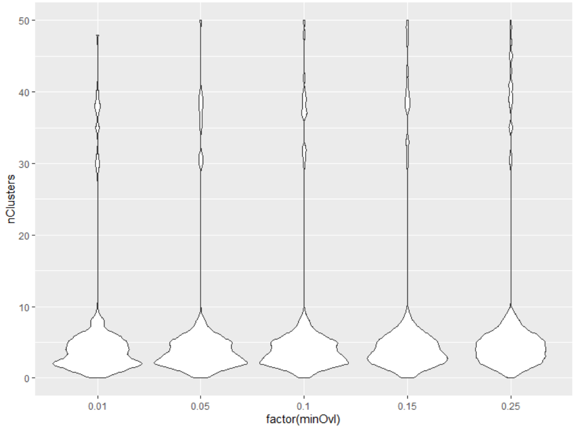


(b)

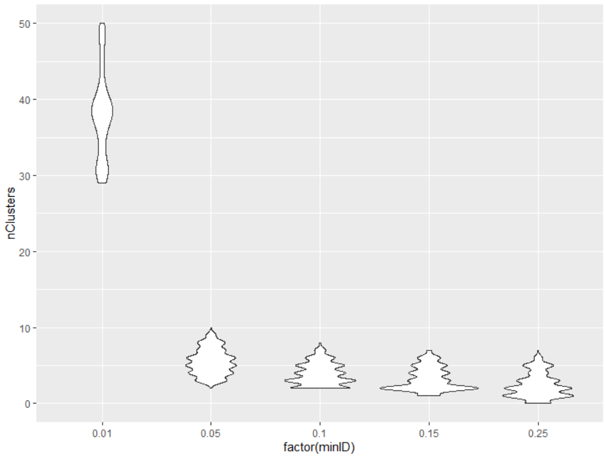


(c)

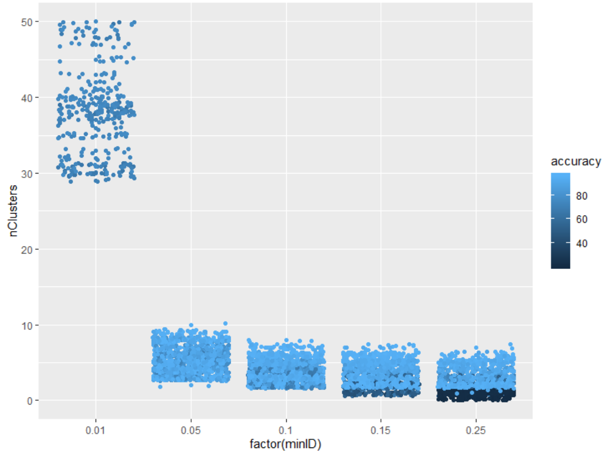

**Supplemental figure 6.** Interaction between ploidy and ID parameter. This graph recalls the one in figure 1 in which we show how different values for the ID parameter lead to differences in accuracy. Her we display these same graphs separated by ploidy, showing that the three ploidies we tested (2n, 3n and 4n) are differently affected by the ID parameter value. As the ploidy increases, the range of values for the ID parameter that lead to accurate results narrows to around 0.05.

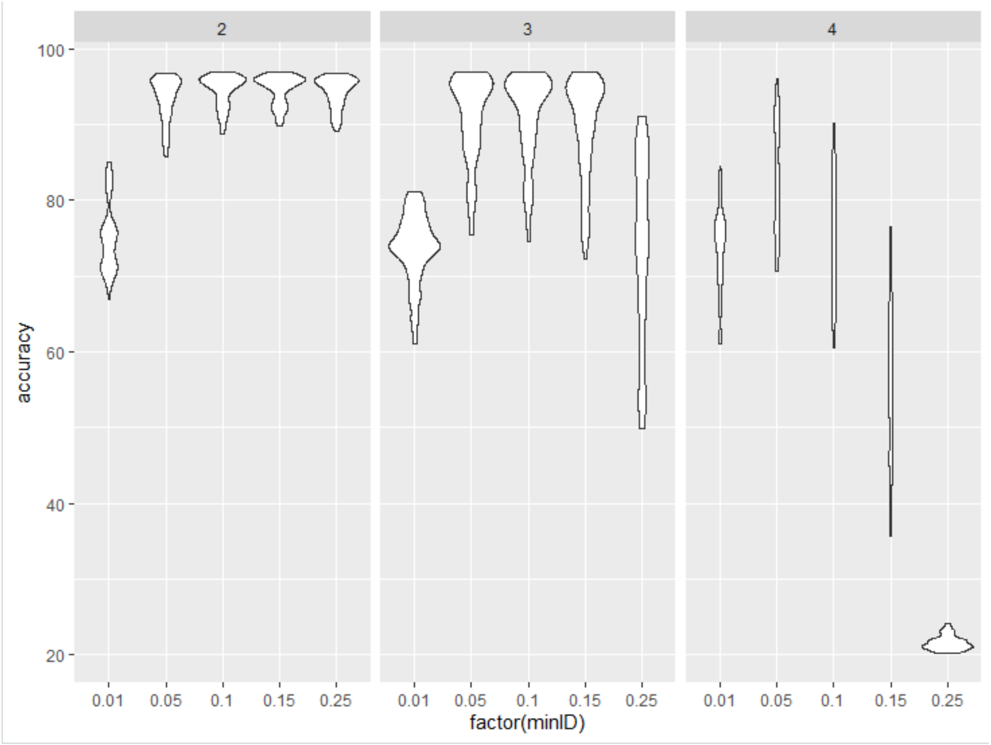

**Supplemental Figure 7.** Effects of coverage on accuracy and contiguity. We compared the results of all 3000 tests performed on 10X datasets to the 3000 tests performed on their 20X counterparts and found that the 20X datasets had consistently more potential, reaching higher accuracy values and better contiguity across ploidy and heterozygosity levels. (a) Here we compare the accuracy distributions for tests of different ploidies and heterozygosity levels at the 10X and 20X coverage levels. We see that the 20X dataset is consistently able to reach higher accuracy levels, an effect which appears to be stronger when the ploidy is higher. (b) Since we are only interested in a high contiguity when it is coupled with a high accuracy, we will not look at a distribution of the number of haplotigs per haplotype across ploidy and heterozygosity levels at the 10X and 20X coverage levels, instead we're focusing on that contiguity metric for the tests using default parameters. We can clearly see a higher number of haplotigs per haplotype for the 10X dataset, with the gap between the 10X and 20X datasets deepening for lower heterozygosity level tests.

(a)

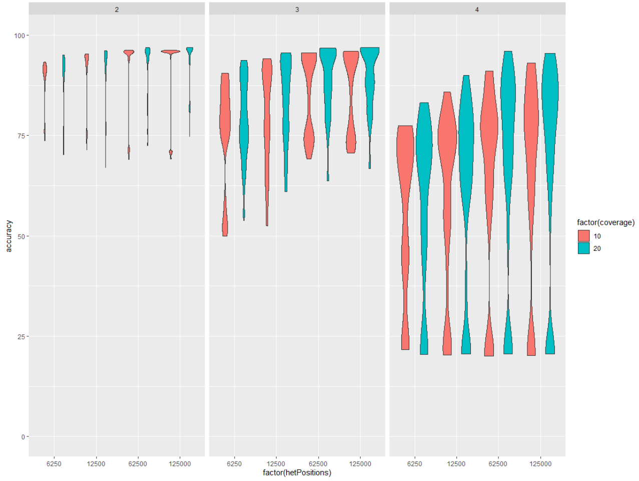


(b)

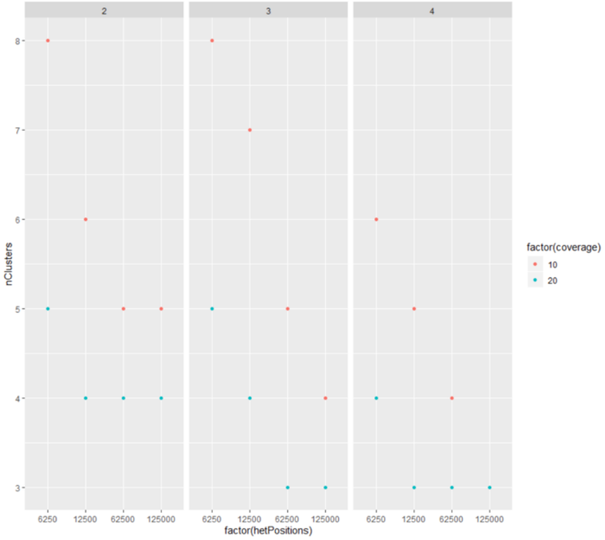

**Supplemental Figure 8.** Effects of including split reads. We ran 3000 tests on all parameter combinations without including split read information and 3000 tests with split read information. These violin plots show the impact of split read information on accuracy and contiguity. (**a**) Based on the violin plots, the results of tests that included split read information display significantly fewer haplotigs, indicative of a higher contiguity. (**b**) The accuracy of results for tests that included split reads is virtually identical to the accuracy of results without split reads.

(a)

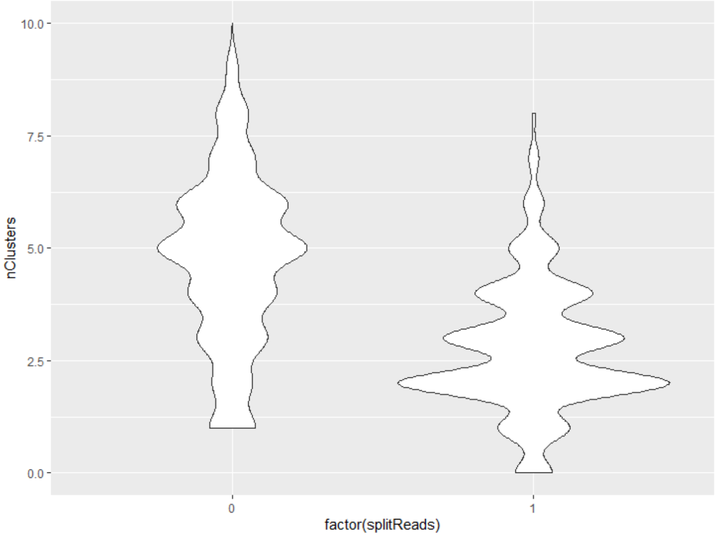


(b)

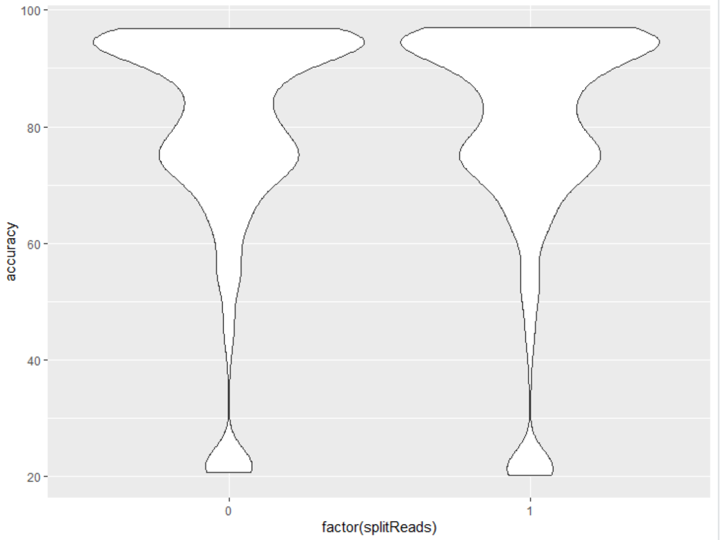
